# Category-selective functional connectivity during episodic encoding and retrieval in younger and older adults

**DOI:** 10.64898/2026.05.29.728795

**Authors:** Sarah A. Monier, Sabina Srokova, Nehal S. Shahanawaz, Michael D. Rugg

**Affiliations:** Center for Vital Longevity and School of Behavioral and Brain Sciences, University of Texas at Dallas 1600 Viceroy Dr. Ste. 800, Dallas, TX 75235; Department of Psychology and Evelyn F. McKnight Brain Institute, University of Arizona 1333 N. Martin Ave. (Building 221), University of Arizona Tucson, AZ 85719

**Keywords:** aging, episodic memory, fMRI, task-based connectivity

## Abstract

Regions within ventral occipito-temporal cortex exhibit category-selective BOLD responses during episodic encoding and retrieval of visual information. How these regions interact with other brain areas during successful encoding and retrieval, and whether these interactions relate to memory performance, remains unclear. The present study examined category-selective functional connectivity using psychophysiological interaction (PPI) analyses in younger and older adults during the encoding and retrieval of word-image associations. Seed regions comprised three scene-selective regions – the parahippocampal place area, medial place area, and occipital place area – and one object-selective region, the lateral occipital complex (LOC). During encoding, scene-selective regions exhibited greater connectivity with posterior occipital and occipitotemporal regions during scene relative to object encoding, whereas the LOC exhibited less extensive connectivity with similar posterior regions during object encoding. During retrieval, both scene- and object-selective regions demonstrated increased connectivity with left lateral prefrontal and parietal cortices during the retrieval of their preferred category. Age differences in scene-selective connectivity were evident at both phases. Moreover, associations between source memory performance and scene-selective connectivity were significant only in younger adults. These findings suggest that scene- and object-selective regions exhibit convergent patterns of functional connectivity during encoding and retrieval which, for scenes, vary with age.

---

Episodic memory refers to the conscious retrieval of unique, personally experienced events (Tulving, 1985). Episodic memories depend on the successful encoding and storage of detailed, context-rich neural representations of such events. Critically, the fidelity of these representations – how precisely and distinctively information is encoded – is a key determinant of subsequent memory success (Rissman & Wagner, 2012; Xue, 2018; Hill et al., 2021).

At encoding, regions within the occipito-temporal cortex exhibit preferential activation to distinct classes of visual stimuli (Koen & Rugg, 2019). For instance, the parahippocampal place area (PPA) responds more robustly to scenes relative to other visual categories (Epstein and Kanwisher, 1998), while the lateral occipital complex (LOC) shows heightened neural activity for objects (Malach et al., 1995; Grill-Spector et al., 2001). Episodic retrieval of visual information is associated with enhanced activity within category-selective neural regions, along with a seemingly content-independent ‘core recollection network’ that includes the hippocampus, medial prefrontal cortex, posterior cingulate, and angular gyrus (Rugg & Vilberg, 2013).

Whereas category-selective activation during encoding and retrieval has been extensively investigated, much less is known about how these regions interact with other brain areas during successful encoding and retrieval, and whether such functional connectivity patterns relate to memory performance. Category-selective responses in the ventral visual pathway are thought to reflect not only regional activity but also the broader distributed networks to which the regions belong to. For example, findings from resting-state fMRI studies suggest that the functional organization of category-selective regions in the ventral visual pathway is reflected in their patterns of connectivity with distributed cortical networks involved in processing category-relevant information (Hutchinson et al., 2014; Stevens et al., 2015; Chen et al., 2017). However, because resting-state connectivity reflects task-independent coupling, these findings do not address whether connectivity between category-selective regions and other cortical areas is modulated by mnemonic demands. Task-based approaches such as psychophysiological interaction (PPI) analysis provide a means to assess how functional connectivity changes as a function of task demands.

Prior task-based functional connectivity studies have largely focused on the hippocampus and surrounding medial temporal lobe (MTL) regions and typically report that successful encoding involves coordinated interactions between the hippocampus and cortical areas involved in perceptual and cognitive control processes (see Palacio & Cardenas, 2019, for a review; Sperling et al., 2003; Ranganath et al., 2005). Evidence also indicates that the encoding of durable episodic memories – those retrieved weeks after the encoding event – are characterized not by greater regional activation, but by increased connectivity between the right hippocampus and posterior perceptual and default mode regions at encoding (Sneve et al., 2015).

Among the studies that examined connectivity at retrieval, King et al. (2015) reported that successful recollection was accompanied by enhanced connectivity between core recollection regions and, for each region, enhanced connectivity with the frontoparietal control network (including the superior parietal and dorsolateral prefrontal cortex). A particularly intriguing aspect of these findings was that regions demonstrating significant functional connectivity with core recollection areas were not limited to those showing elevated activity during successful retrieval but extended to other neocortical areas, including visual cortex.

In another study (Wais et al., 2017), high-fidelity recognition memory was associated with increased connectivity between medial temporal and frontoparietal regions, whereas no such connectivity changes were identified for low-fidelity recognition. Using a graph-theoretical approach, Schedlbauer et al. (2014) reported that accurate retrieval (i.e., correct vs. incorrect spatiotemporal judgements) elicited widespread increases in connectivity among hippocampal, prefrontal, parietal, and visual regions, and that overall connectivity strength predicted memory performance (see also King et al., 2015). Together, these findings suggest that successful episodic memory relies on coordinated activity between multiple regions.

While prior work has underscored the importance of examining distributed functional connectivity changes during memory encoding and retrieval, an open question remains whether category-selective regions, such as scene- and object-selective visual cortices, exhibit distinct connectivity patterns that vary with the content of the encoded or retrieved information. In one of the few studies to investigate task-dependent connectivity of category-selective regions, Summerfield et al. (2006) employed a ‘state-related’ connectivity approach to examine whether functional connectivity between prefrontal cortex and the fusiform face area (FFA) and parahippocampal place area (PPA) varied as a function of encoding success. They found that stronger connectivity between these regions and the left dorsolateral prefrontal cortex (DLPFC) was associated with successful associative encoding, consistent with the proposal that top-down control mechanisms interact with category-specific perceptual regions to guide the selection of task-relevant information.

A further question concerns whether task-dependent connectivity patterns vary with age (see Cansino et al., 2022, for a review). In most of these studies, age-related differences in connectivity were examined with respect to medial temporal regions, particularly connectivity between the hippocampus and neocortical regions outside of the medial temporal lobe. Notably, age differences have been reported not only in the strength of connectivity but also in its relationship with memory performance (Ankudowich et al., 2019; Monge et al., 2018), although one study found that age differences in recollection-related connectivity were no longer significant after accounting for individual differences in memory performance (King et al., 2018). To our knowledge, however, no study has examined age-related differences in the functional connectivity of category-selective regions and how these effects relate to individual differences in memory performance.

The present study extends prior work by examining how category-selective cortical regions functionally connect with other regions during content-selective encoding and retrieval in younger and older adults. Participants underwent fMRI while performing an associative memory task, and PPI was used to quantify task-dependent changes in functional connectivity between scene- and object-selective regions and the rest of the brain. The primary goals were to determine whether successful encoding and retrieval are associated with the modulation of category-specific connectivity patterns, and whether these patterns differ across age groups. A further goal was to assess whether connectivity changes at encoding and retrieval are associated with individual differences in memory performance, and whether these relationships vary with age.

## Materials and Methods

The experimental design, procedures, and behavioral results have been described elsewhere, along with eye movement data from the study phase (Srokova et al., 2025). Note that the participant sample in Srokova et al., 2025 does not completely overlap with that employed in the current study. For the convenience of the reader, we provide a detailed description of the experimental procedures and behavioral results relevant to the analyses of the encoding- and retrieval-related functional connectivity effects that are the focus of the current paper.

### Participants

26 young (18-30 yrs) and 32 older adults (65-75 yrs) were recruited from the University of Texas at Dallas and surrounding metropolitan Dallas communities. All participants were right-handed, had normal or corrected-to-normal vision, and were fluent in English before age five. Participants were compensated at $30 per hour and up to $30 for travel. All participants provided written informed consent in accordance with the University of Texas at Dallas Institutional Review Board, which approved the study (approval number IRB-19-124). Exclusion criteria included any history of neurological or psychiatric disorders, substance abuse, diabetes, and current or recent use of medications affecting the CNS.

Prior to MRI scanning, all participants completed neuropsychological testing. Standard criteria were applied to exclude potential cases of early cognitive decline or mild cognitive impairment (see the ‘Neuropsychological testing’ section below). Of the participants who met the inclusion criteria, two young and five older adults were excluded from all analyses. Exclusions were due to an incidental MRI finding (n=1), technical malfunction during scanning (n = 1), falling asleep during MRI (n = 1), using the incorrect hand for responses (n = 1), and near-chance source memory performance (pSR < 0.1; n = 3). The final sample comprised 24 young and 27 older adults.

### Neuropsychological testing

Participants underwent a neuropsychological evaluation on a day prior to the MRI session. The test battery comprised the Mini-Mental State Examination (MMSE), the California Verbal Learning Test II (CVLT; Delis et al., 2000), the Wechsler Logical Memory Tests 1 and 2 (Wechsler, 2009), the Symbol Digit Modalities Test (SDMT; Smith, 1982), the Trail Making Tests A and B (Reitan and Wolfson, 1985), the F-A-S subtest of the Neurosensory Center Comprehensive Evaluation for Aphasia (Spreen and Benton, 1977), the Digit Span Forward and Backward subtests of the revised Wechsler Adult Intelligence Scale (WAIS; Wechsler, 1981), the Category Fluency test (Benton, 1968), Raven’s Progressive Matrices List I (Raven et al., 2000), and the Test of Premorbid Functioning (TOPF; Wechsler, 2011).

Additionally, participants’ visual acuity was measured using the Early Treatment Diabetic Retinopathy Study (ETDRS) charts with the logMAR metric, with participants wearing prescribed corrective lenses when applicable. Potential participants were excluded from the MRI session if they scored: > 1.5 SD below the age-appropriate norm on two or more non-memory assessments, > 1.5 SD below the age-appropriate norm on at least one memory-based test, or < 26 on the MMSE.

### Experimental materials

The critical experimental stimuli comprised 192 concrete nouns and 128 colored images depicting 64 scenes (32 indoor/32 outdoor) and 64 objects (32 manmade/32 natural). Additional stimuli included 16 words and 16 images (8 scenes, 8 objects) reserved for practice trials or as fillers at the beginning of each scanner run. For the study phase, critical stimuli consisted of 128 word-image pairs (evenly distributed across image categories) interspersed with 32 null trials (a white fixation cross lasting 4 seconds). For the test phase, critical stimuli consisted of 128 previously studied words, 64 unstudied words, and 48 null trials. The critical stimuli were used to create a total of 27 stimulus lists. 24 of these were assigned to yoked pairs of younger and older adults, and the remaining three lists were assigned to three older adults. The order of critical items was pseudorandomized such that no more than three successive trials of the same category were presented, and no more than two null trials were presented consecutively. Experimental stimuli were presented using PsychoPy v2021.1.3 (Peirce et al., 2019). Practice lists for the study and test phases were created using a separate pool of stimuli.

### Experimental procedure

Participants completed two study-test cycles of an associative memory task inside the MRI scanner (see Figure 1). Each cycle consisted of two consecutive study runs (5 minutes each) followed by two consecutive test runs (10 minutes each). Each trial of the study phase commenced with the presentation of a red fixation cross (500 ms) at the center of the screen, followed by the presentation of a word (1000 ms). An image depicting either a scene or an object then appeared (4000 ms), followed by a white fixation cross displayed for 1000 ms. On scene trials, participants were instructed to imagine the object denoted by the word moving around or interacting within the scene. On object trials, participants imagined the object denoted by the word interacting with the displayed object. Participants had 5 seconds from the onset of the image to rate the vividness of their imagined scenario on a three-point scale: ‘Not Vivid’ (index finger), ‘Somewhat Vivid’ (middle finger), ‘Very Vivid’ (ring finger). Responses were made on a scanner-compatible button box using the right hand.

**Figure 1.**
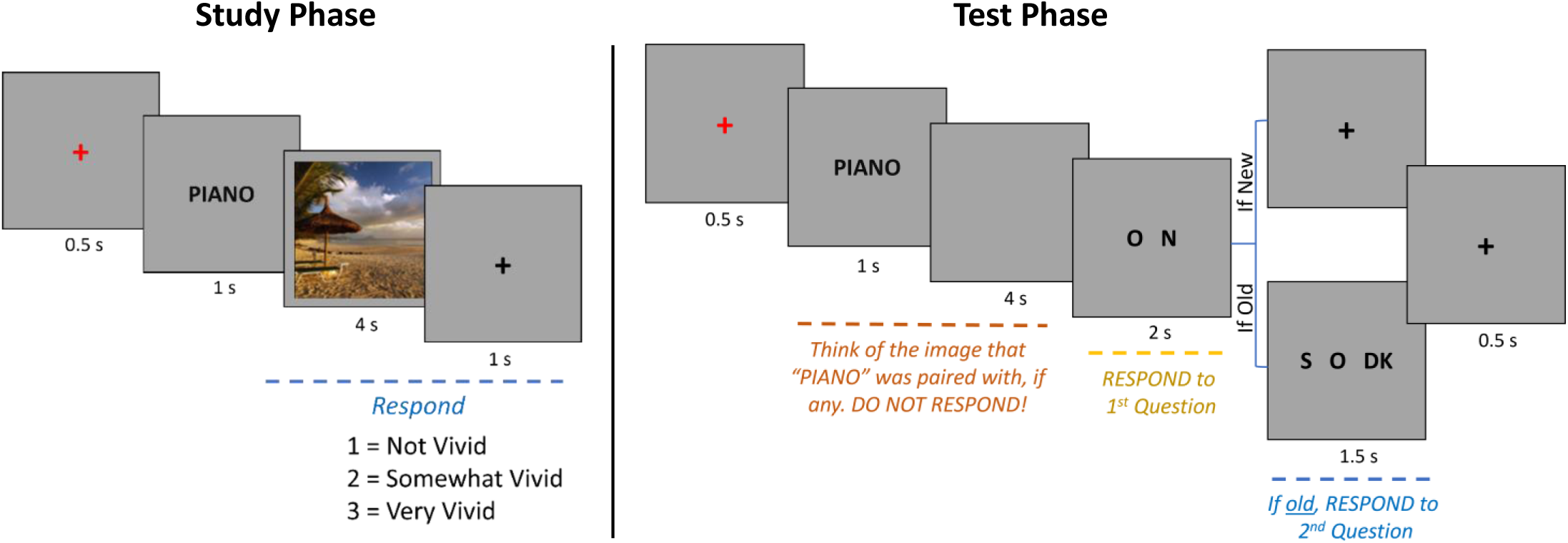
Schematic illustration of the study and test phases. During the study phase, participants were presented with words paired with images of objects or scenes and were instructed to imagine a scenario in which the object denoted by the word interacted with the image. In the test phase, participants were shown a mix of old and new words. Each word was followed by a blank screen for 4 seconds, during which the task was to recall the associated image that had been paired with the word at study. Following this imagery period, participants indicated whether the word was old or new, and if old, whether it had been paired with a scene or an object. **Alt text:** Illustration of the study and test phases of the episodic memory task depicting word-image study trials and test trials requiring old/new and source memory responses.

Test trials began with a red fixation cross (500 ms), followed by the presentation of a word for 1000 ms. Subsequently, a blank screen was displayed for 4000 ms, during which participants were instructed to imagine the image that had been paired with the word during study. Following this visualization period, participants had 2000 ms to indicate whether the word was ‘Old’ (previously studied) or ‘New’ (not studied), using the index and middle fingers of the right hand. Finger-mapping for the ‘Old’ and ‘New’ responses was counterbalanced across participants. For words judged as ‘Old’, an additional 2000 ms was allotted to make a source memory judgement indicating whether the word had been paired with a scene or an object. A third ‘Don’t know’ option was available to discourage guessing. Responses were made using a scanner-compatible button box with the index, middle, and ring fingers of the right hand, and were counterbalanced across participants.

### MRI data acquisition and preprocessing

Functional and structural MRI data were acquired at the Sammons BrainHealth Imaging Center at the University of Texas at Dallas using a Siemens Prisma 3T scanner equipped with a 32-channel head coil. A T1-weighted anatomical image was acquired with a 3D MPRAGE pulse sequence (FOV = 256 x 224, voxel size 1 x 1 x 1 mm, 160 slices, sagittal acquisition). Functional scans were acquired using a T2*-weighted BOLD echoplanar imaging (EPI) sequence with a multiband factor of 3 (flip angle = 70° FOV = 220 × 220 mm; voxel size = 2 × 2 × 2 mm; TR = 1.52 ms; TE = 30 ms; 66 slices). A double-echo fieldmap sequence was collected immediately after the final run of the study phase (TE 1 = 4.92 ms, TE 2 = 7.38 ms, TR = 1.52 ms, FOV = 220 x 220 mms, flip angle = 70°) immediately following the last run of the study phase. The acquisition produced two magnitude images (one for each echo), and a pre-subtracted phase image, calculated as the differences between the phases acquired at each echo.

The data were preprocessed and analyzed using Statistical Parametric Mapping (SPM12; http://www.fil.ion.ucl.ac.uk) and custom MATLAB code (MathWorks). First, voxel displacement maps were generated to correct for distortions caused by magnetic field inhomogeneities. Next, the data were realigned to the mean EPI image and a dynamic correction was applied using the displacement maps.

The images were then reoriented to align with the anatomical image orientation and spatially normalized to an age-unbiased sample-specific EPI template. Finally, the data were smoothed with a 5-mm full-width half-maximum (FWHM) Gaussian kernel.

### fMRI Data Analysis

#### Whole-brain univariate analyses

Separate statistical models were constructed for the encoding and retrieval phases. For each phase, a whole brain analysis of the fMRI data was conducted using a two-stage univariate general linear model (GLM) approach.

#### Encoding-related univariate effects

First-level GLMs were constructed for each participant’s study data. Study trials were binned into four events of interest based on subsequent memory performance at test: scene source correct, object source correct, scene source incorrect, and object source incorrect. Trials were classified as source correct if the associated image category (scene or object) was correctly identified at retrieval. Trials receiving an incorrect source judgment or a ‘Don’t know’ source response were categorized as source incorrect. Each event was modeled with a 4-second boxcar function, corresponding to the duration that the image was presented at study, and convolved with SPM’s canonical hemodynamic response function (HRF). Events of no interest included filler trials, trials that did not receive a vividness judgment, and trials that received a response before 500 ms post-image onset. Other covariates of no interest included the mean signal from each scanner run, 6 motion regressors representing rigid-body translation and rotation, and spike covariates that removed volumes with transient displacements greater than 1 mm or 1° in any direction.

In the second stage, participant-specific parameter estimates derived from the first-level GLM were submitted to a 2 (age group) x 4 (item type) mixed factorial ANOVA. Results of the group-level analysis were considered statistically significant if they exceeded a voxel wise threshold of p < .001 with cluster-level FWE correction at p < .05. Scene-selective encoding effects were operationalized by contrasting neural activity for study trials later associated with a correct scene source judgment versus correct object source judgment, and vice versa for object-selective encoding effects.

#### Retrieval-related univariate effects

For the test data, the first level again involved constructing separate GLMs for each participant. Test trials were binned into 5 events of interest: (1) scene trials associated with a correct source memory judgment (scene source correct), (2) object trials associated with a correct source memory judgment (object source correct), (3) scene trials receiving an incorrect or a ‘Don’t know’ source memory response (scene source incorrect), (4) object trials receiving an incorrect or a ‘Don’t know’ source memory response (object source incorrect), and (5) unstudied items correctly identified as ‘new’ (correct rejections). Each event of interest was modeled with a 5-second boxcar function, corresponding to the duration of the word onset and the imagery period during which participants visualized the image that had been associated with the word at study (see Figure 1). Boxcar regressors were convolved with SPM’s canonical HRF. Events of no interest included filler trials, false alarms, and trials with no response. Additional covariates of no interest included the mean signal of each scan session, six rigid-body motion parameters and spike regressors coding for volumes with a transient displacement of > 1 mm or > 1 degree.

In the second stage, participant-specific parameter estimates derived from the first-level GLMs were carried over to a 2 (age group) x 5 (item type) mixed factorial ANOVA. Statistical significance was defined using the same criteria as for the study data analysis (see above). Category-selective retrieval effects were operationalized as greater neural activity for source correct trials of the preferred category versus source correct trials of the non-preferred category. This contrast was employed because a sizeable subset of participants had too few source incorrect trials to permit reliable estimation of recollection-related effects. Accordingly, analyses were restricted to source correct trials to maximize sample size and hence statistical power.

### Psychophysiological Interaction Analyses

To examine modulations of functional connectivity (FC) associated with successful category-selective encoding and retrieval, Psychophysiological Interaction (PPI) analyses were performed using multiple seed regions (Friston et al., 1997).

#### Selection of category-selective seed regions

Category-selective seeds in the present analyses included three scene-selective regions- the parahippocampal place area (PPA), medial place area (MPA; also sometimes referred to as the retrosplenial complex), and occipital place area (OPA) - and one object-selective region, the lateral occipital complex (LOC). Each of these regions is illustrated in Figure 3. The regions were selected based on strong *a priori* evidence of their functional specialization for scene and object processing, respectively. In every case, the seeds fell within regions demonstrating category-selective encoding and retrieval effects as defined above.

To avoid circularity, seed regions were defined based on a prior independent localizer dataset following the same procedures as those employed by Srokova et al. (2021). Briefly, localizer data were analyzed in two stages prior to seed definition. First-level GLMs were estimated for each participant, modeling three events of interest: scene, object, and scrambled blocks. Each block was modeled with a 12-second boxcar function and convolved with a canonical hemodynamic response function (HRF) and its derivatives. Nuisance regressors included six motion parameters, session mean signals, and spike regressors for volumes exceeding 1 mm translation or 1° rotation. Subject-level estimates were entered into a second level 2 (age group: younger, older) x 3 (stimulus type: scene, object, scrambled) mixed-effects ANOVA.

The PPA, MPA, and OPA seeds were derived using a conjunction of group-level scene > object + scrambled image contrasts. The contrasts were masked using labels from the Neuromorphometrics atlas available in SPM12 and restricted the PPA to the parahippocampal/fusiform cortex and the OPA to the inferior and middle occipital gyri. Given that Neuromorphometrics does not offer a precise anatomical mask for the MPA, this region was defined by applying an inclusive mask to the contrast using results from a Neurosynth database search for the term “retrosplenial” (conducted in August 2019; results were FDR-corrected at p < .00001; Yarkoni et al., 2011). The LOC seed was defined using the group-level contrast of object > scene + scrambled image, and the resulting cluster was inclusively masked using atlas-defined labels for the inferior and middle occipital gyri to anatomically constrain the region of interest (ROI). The number of voxels contained within each seed region was 336 and 421 (left and right PPA, respectively), 254 and 400 (left and right MPA), 531 and 456 (left and right OPA) and 818 and 778 (left and right LOC).

Preliminary connectivity analyses employing hemisphere-specific seed regions yielded results highly consistent with those obtained using bilateral seeds. Accordingly, the left and right hemisphere seeds were combined to form a single seed region for the analyses reported below. However, object-selective connectivity at retrieval was assessed using only the left LOC as a seed, as category-selective univariate retrieval effects were observed only in this region. Because of the poor overlap between the localizer-defined LOC mask and object-selective univariate retrieval effects (see Figure 3), we defined a separate retrieval-related left LOC seed using all voxels demonstrating a significant object-selective univariate retrieval effect in the whole-brain analyses (height threshold: p < .001, uncorrected; cluster-level FWE-corrected p < .05). The resulting cluster comprised 102 voxels.

#### First-level PPI models

PPI analyses were conducted separately for the study and test phases and for each seed region of interest. For each seed, the PPI model incorporated three regressors in the subject-level GLMs: a physiological regressor, a psychological regressor, and the PPI interaction term. The first eigenvariate of the time series from all voxels within a seed served as the physiological regressor. For both the encoding and retrieval phases, the psychological regressor was defined by a task vector that represented category-selective source memory performance (coded as 1 for source-correct trials of the preferred category, -1 for source-correct trials of the non-preferred category, and 0 for all other trials). To create the PPI interaction term, the extracted time courses were deconvolved with the canonical HRF and multiplied by the task vector. The product from this step was then reconvolved with the HRF. The PPI interaction term was entered into the first-level GLM as the regressor of interest.

#### Second-level PPI analyses

First-level parameter estimates were carried forward to second-level random-effects analyses. Separate contrasts were specified to assess the main effect of connectivity change, group-specific effects, and age-related differences in PPI (i.e., age group x PPI interaction). For each significant seed-target connectivity effect (using the same thresholds described previously; see ‘Whole-brain univariate analyses’), PPI parameter estimates were extracted from 5 mm radius spheres centered on the peak voxels of the significant PPI interaction term for each participant.

Given the difficulty in interpreting negative PPI effects, the present analyses focused exclusively on relative increases in functional connectivity associated with the contrast of interest. Accordingly, only increased functional connectivity effects are reported and interpreted.

### Age differences in functional connectivity

To assess whether age modulated any of the category-selective functional connectivity effects at encoding and retrieval, participant-specific parameter estimates (see above) were subjected to mixed-design ANOVAs for each seed, with age group (younger, older) as a between-subjects factor and target region as a within-subjects factor. Greenhouse-Geisser corrections were applied when the assumption of sphericity was violated, and effects were considered significant at p < .05. In cases where extreme values were identified, analyses were conducted both before and after winsorization of connectivity estimates. Winsorization involved replacing values below the 5^th^ percentile with the 5^th^ percentile value, and values above the 95^th^ percentile with the 95^th^ percentile value.

### Relationships between behavioral performance and connectivity metrics

To examine whether category-selective connectivity differences were associated with memory performance, separate multiple regression analyses were conducted to predict item recognition accuracy (collapsed across image categories) and source memory performance from the variables of PPI-derived connectivity parameter estimates, age group, and their interaction. In cases where the interaction term was non-significant, the regression models were re-run after removing the term. As described above (see ‘Age differences in functional connectivity’), analyses were performed both prior to and following winsorization of connectivity estimates.

## Results

### Demographic information and neuropsychological test battery results

Demographic information and neuropsychological test performance for young and older participants are summarized in Table 1. Older adults had more years of education and scored lower on the CVLT Short Delay Free Recall, WMS Logical Memory II, SDMT, Trails A and B, and TOPF relative to young adults.

**Table 1.**
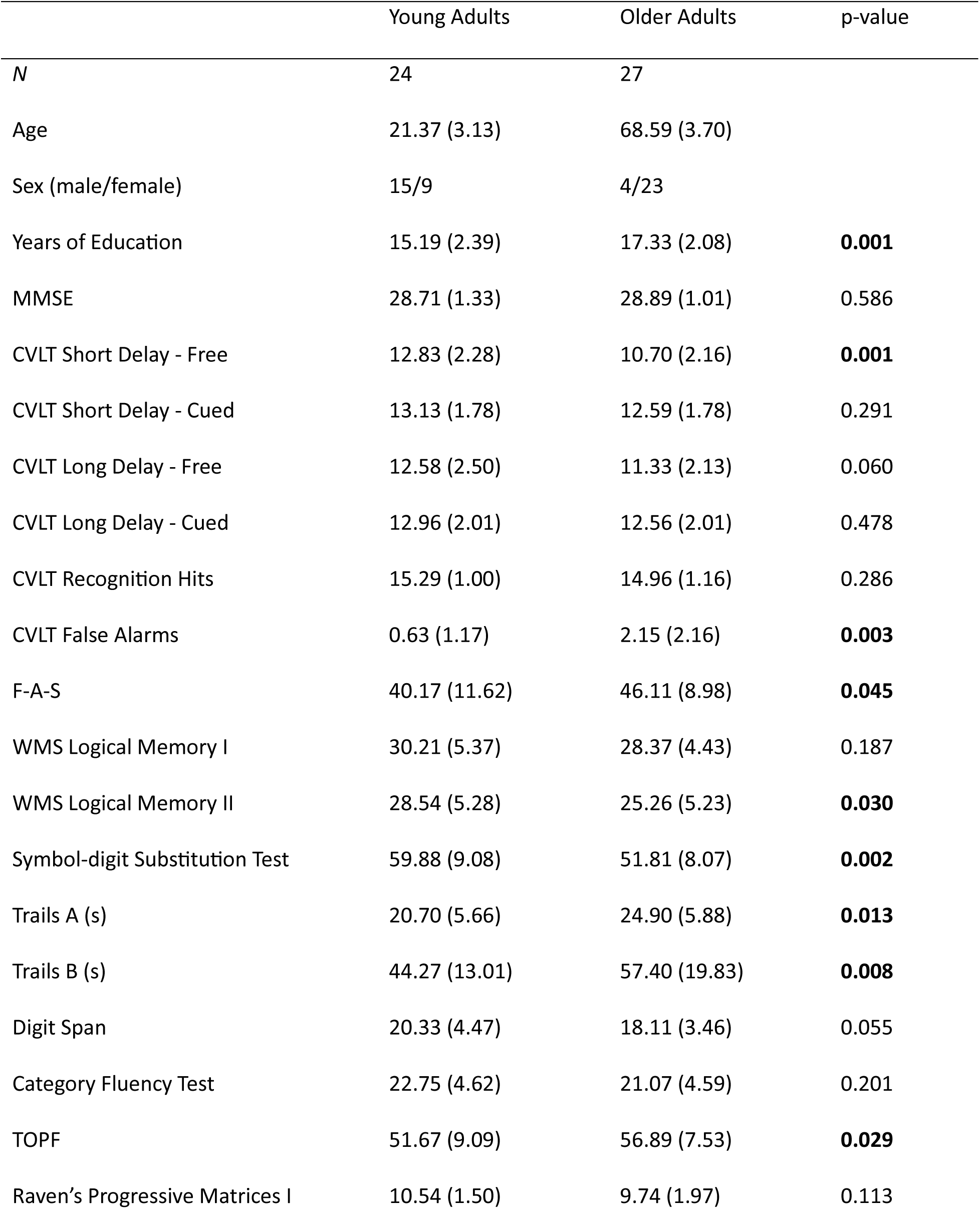
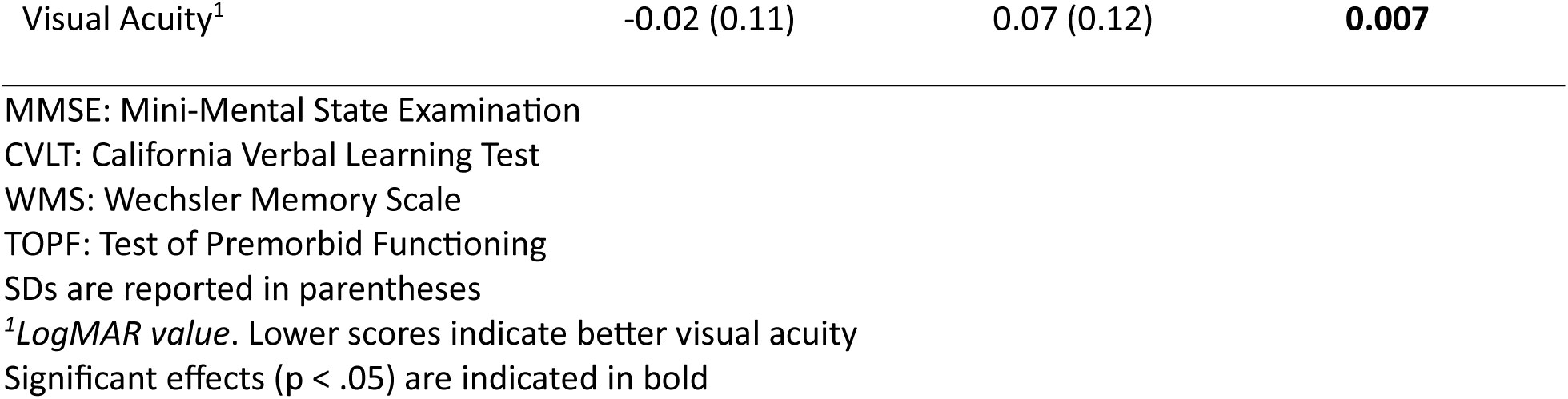
Participant demographic information and neuropsychological test battery scores in younger and older adults.

|  | Young Adults | Older Adults | p-value |
| --- | --- | --- | --- |
| *N* | 24 | 27 |  |
| Age | 21.37 (3.13) | 68.59 (3.70) |  |
| Sex (male/female) | 15/9 | 4/23 |  |
| Years of Education | 15.19 (2.39) | 17.33 (2.08) | **0.001** |
| MMSE | 28.71 (1.33) | 28.89 (1.01) | 0.586 |
| CVLT Short Delay - Free | 12.83 (2.28) | 10.70 (2.16) | **0.001** |
| CVLT Short Delay - Cued | 13.13 (1.78) | 12.59 (1.78) | 0.291 |
| CVLT Long Delay - Free | 12.58 (2.50) | 11.33 (2.13) | 0.060 |
| CVLT Long Delay - Cued | 12.96 (2.01) | 12.56 (2.01) | 0.478 |
| CVLT Recognition Hits | 15.29 (1.00) | 14.96 (1.16) | 0.286 |
| CVLT False Alarms | 0.63 (1.17) | 2.15 (2.16) | **0.003** |
| F-A-S | 40.17 (11.62) | 46.11 (8.98) | **0.045** |
| WMS Logical Memory I | 30.21 (5.37) | 28.37 (4.43) | 0.187 |
| WMS Logical Memory II | 28.54 (5.28) | 25.26 (5.23) | **0.030** |
| Symbol-digit Substitution Test | 59.88 (9.08) | 51.81 (8.07) | **0.002** |
| Trails A (s) | 20.70 (5.66) | 24.90 (5.88) | **0.013** |
| Trails B (s) | 44.27 (13.01) | 57.40 (19.83) | **0.008** |
| Digit Span | 20.33 (4.47) | 18.11 (3.46) | 0.055 |
| Category Fluency Test | 22.75 (4.62) | 21.07 (4.59) | 0.201 |
| TOPF | 51.67 (9.09) | 56.89 (7.53) | **0.029** |
| Raven's Progressive Matrices I | 10.54 (1.50) | 9.74 (1.97) | 0.113 |
| Visual Acuity<sup>1</sup> | -0.02 (0.11) | 0.07 (0.12) | **0.007** |

MMSE: Mini-Mental State Examination
CVLT: California Verbal Learning Test
WMS: Wechsler Memory Scale
TOPF: Test of Premorbid Functioning
SDs are reported in parentheses
<sup>1</sup>*LogMAR value*. Lower scores indicate better visual acuity
Significant effects ( $p < .05$ ) are indicated in bold

### Item recognition and source memory performance

Means and standard deviations of the memory performance measures are reported in Table 2.

**Table 2.**
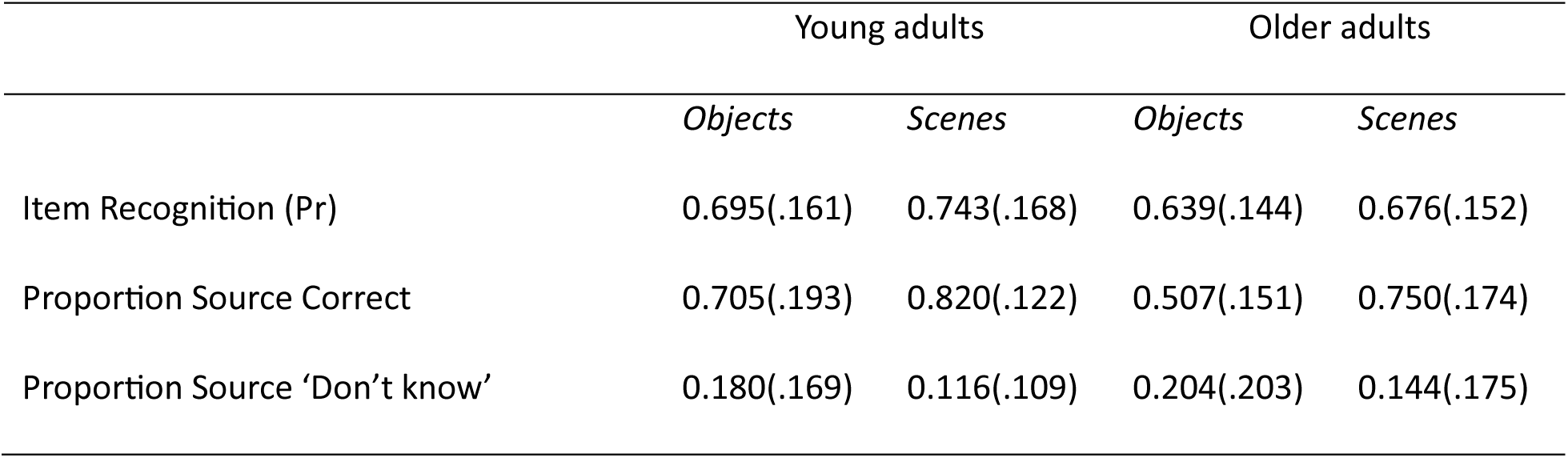
Means (SD) for item and source memory performance.

Item recognition accuracy (Pr) was calculated separately for each image category as the difference between the item hit rate (proportion of correctly recognized old items) and the false alarm rate (the proportion of new items incorrectly identified as “old”). A mixed-design ANOVA revealed a significant main effect of image category on Pr (F(1,49) = 20.054, p < .001, partial η^2^ = 0.290), with higher Pr for scenes (mean [SD] = 0.707 [0.162] than objects (mean [SD] = 0.666 [0.153]. The main effect of age group was not significant (F(1,49) = 2.086, p = 0.155, partial η^2^ = 0.041), and nor was the interaction between age group and image category (F(1,49) = 0.334, p = 0.566, partial η^2^ = 0.007), Source memory performance, operationalized as the probability of source recollection (*pSR*), was estimated using a modified single high-threshold model that accounts for guesses (Snodgrass and Corwin, 1988):

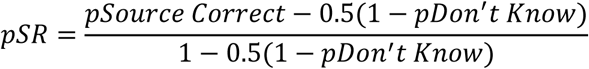

“pSource Correct” and “pDon’t Know” in the formula refer to the proportion of correct “old” trials that received an accurate or ‘Don’t know’ source memory judgement, respectively. An independent samples t-test revealed that *pSR* was significantly lower in older (Mean [SD] = .391 [.185]) than in younger adults (Mean [SD] = .608 [.215]), t(49) = 3.879, p < .001 (see Figure 2).

**Figure 2.**
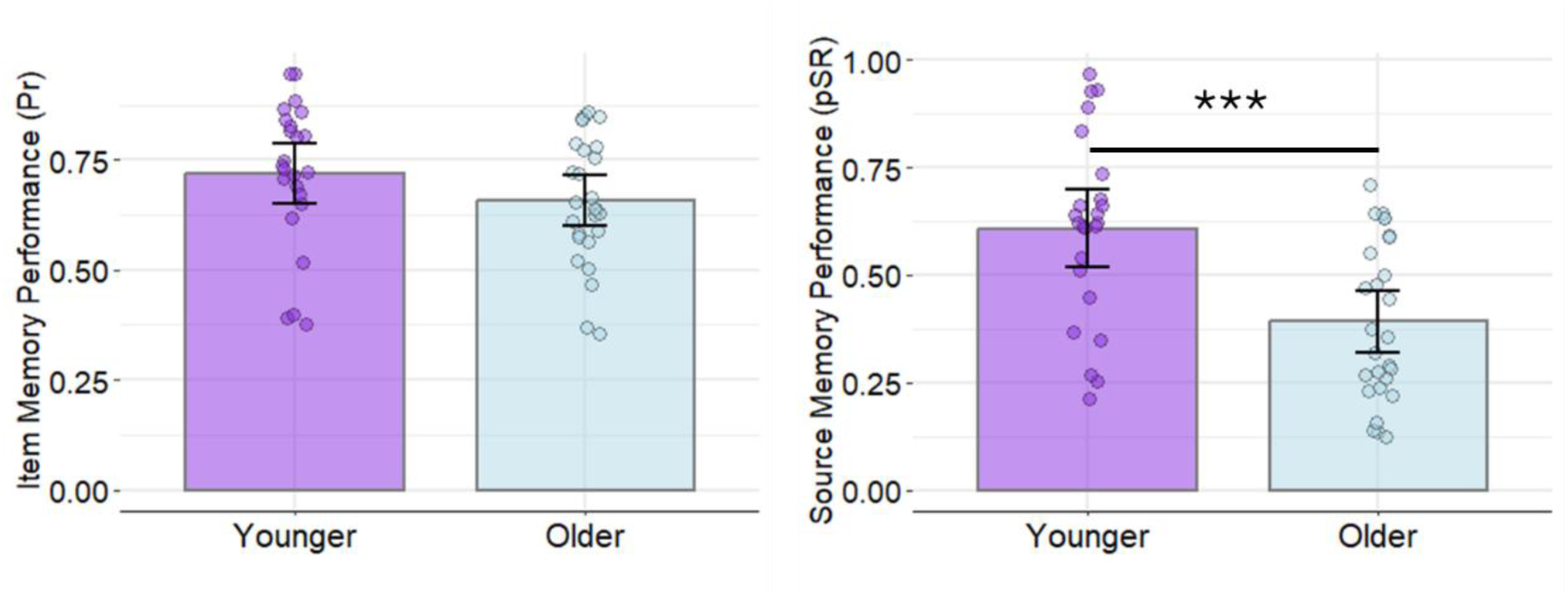
Source and item memory performance in younger and older adults. Item memory performance (Pr) is collapsed across image category. Error bars represent 95% confidence intervals. ***p < .001. **Alt text:** Bar graphs comparing item and source memory performance in younger and older adults. Younger adults showed higher source memory performance than older adults, whereas item memory performance was equivalent across the age groups.

In addition, we examined source accuracy (pSource correct) as a function of image category using a 2 (age group: younger, older) x 2 (category: scenes, objects) mixed-design ANOVA. The ANOVA revealed a significant main effect of category, F(1,49) = 110.217, p < .001, partial η^2^ = 0.692, with scenes receiving a higher proportion of source correct responses (Mean [SD] = .783 [.154]) than objects (Mean [SD] = .600 [.198]). A significant main effect of age group also emerged, F(1,49) = 10.107, p = .003, partial η^2^ = 0.171, indicating that younger adults (Mean [SD] = .763 [.151]) had a greater proportion of source correct responses than older adults (Mean [SD] = .628 [.150]). This analysis also revealed a significant age group x category interaction, F(1,49) = 14.368, p < .001, partial η^2^ = 0.227. Independent t-tests revealed that younger adults had a higher proportion of source correct responses for objects than older adults (t(49) = 4.111, p < .001) but no corresponding age difference was observed for scenes (t(49) = 1.642, p = .107).

### Whole-brain univariate category-selective encoding and retrieval effects

Anatomical labels for significant clusters and peak coordinates were assigned based on the Automated Anatomical Labeling (AAL) atlas.

#### Encoding

As illustrated in Figure 3, scene-selective effects at study were observed in bilateral parahippocampal/medial fusiform cortex (corresponding to the parahippocampal place area [PPA]), ventral posterior cingulate/retrosplenial cortex (medial place area [MPA]), and the occipital place area (OPA), along with several additional cortical regions (see Supplementary Table 1). As also illustrated in Figure 3, object-selective effects at study were identified in bilateral lateral occipital complex (LOC), among other regions (see Supplementary Table 1 for a full list).

**Figure 3.**
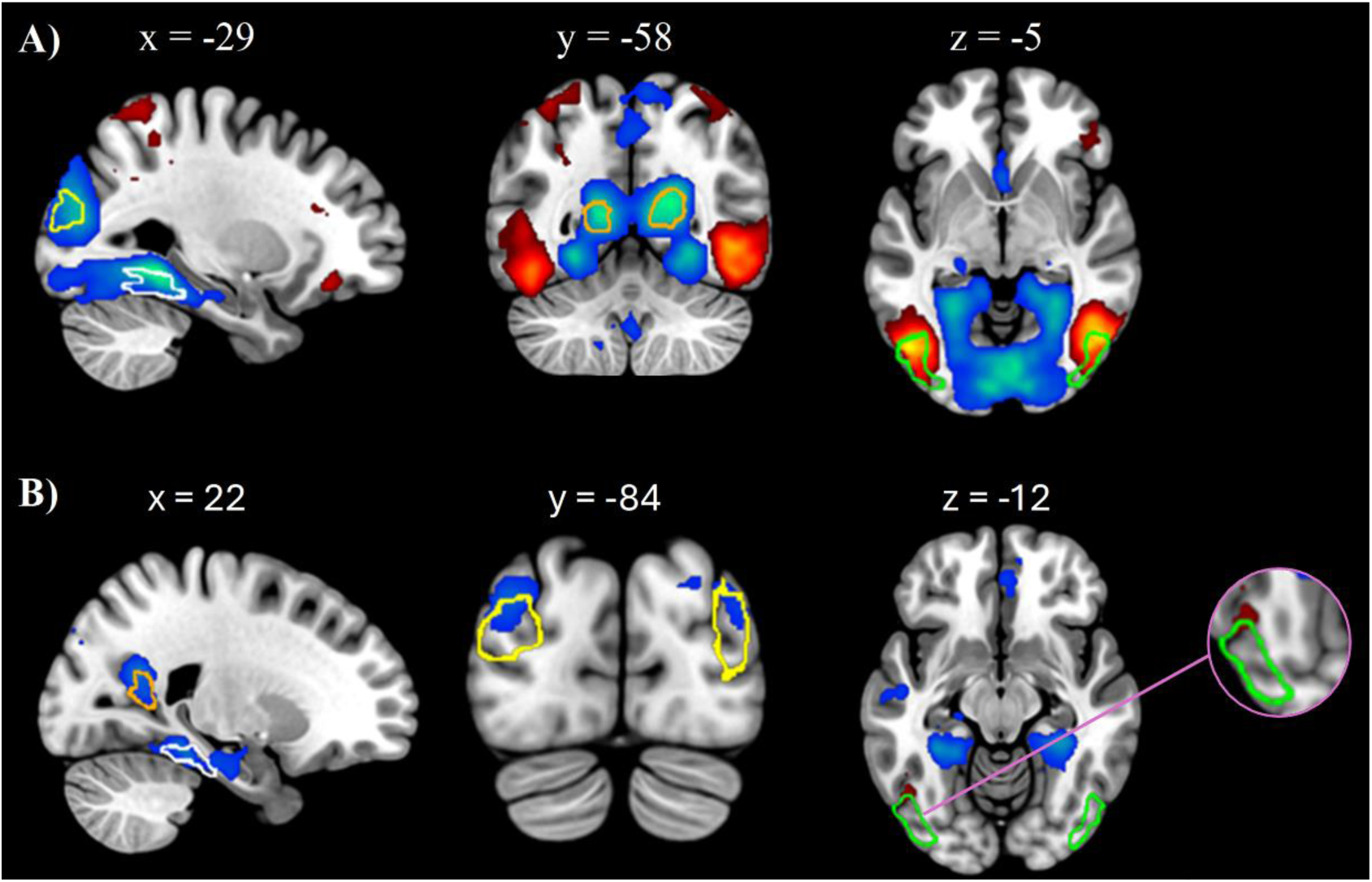
Whole-brain Univariate Effects. (A) Category-selective encoding effects, operationalized as greater activation for scene versus object items later associated with a correct source response (cool colors), and the reverse contrast (warm colors). (B) Category-selective retrieval effects, operationalized as scene source correct > object source correct (cool colors), and the reverse contrast (warm colors). Clusters are displayed at height threshold: p < .001 (uncorrected) with cluster-level FWE correction (p < .05). Scene-and object-selective ROI masks derived from an independent localizer dataset are overlaid on the encoding and retrieval effects to illustrate localization. Color coding of ROI masks: yellow = OPA; orange = MPA; white = PPA; green = LOC. Results are displayed on the Montreal Neurological Institute (MNI) brain using MRIcroGL (https://www.nitrc.org/projects/mricrogl; Brett et al., 2001). **Alt text:** Brain images illustrating whole-brain category-selective activity during encoding and retrieval. Scene- and object-selective region of interest masks are overlaid to illustrate overlap with activation effects.

#### Retrieval

At retrieval, scene-selective effects were again observed in bilateral PPA, MPA, and OPA, along with other cortical regions (Figure 3; also see Supplementary Table 2). In contrast, object-selective effects were identified in the left LOC only (Figure 3).

### Category-selective connectivity effects at encoding

#### Scene-selective effects

As illustrated in Figure 4 and reported in Table 3 (also see Supplementary Table 3 for a more detailed list including subpeak coordinates), scene-selective seeds (MPA, PPA, and OPA) exhibited robust increases in functional connectivity with posterior occipital and occipito-temporal regions, including the fusiform gyrus and the middle and inferior occipital gyri, during the encoding of subsequently remembered scenes relative to subsequently remembered objects. Additionally, the MPA demonstrated increased connectivity with left lateral prefrontal cortex, particularly the inferior frontal gyrus.

**Figure 4.**
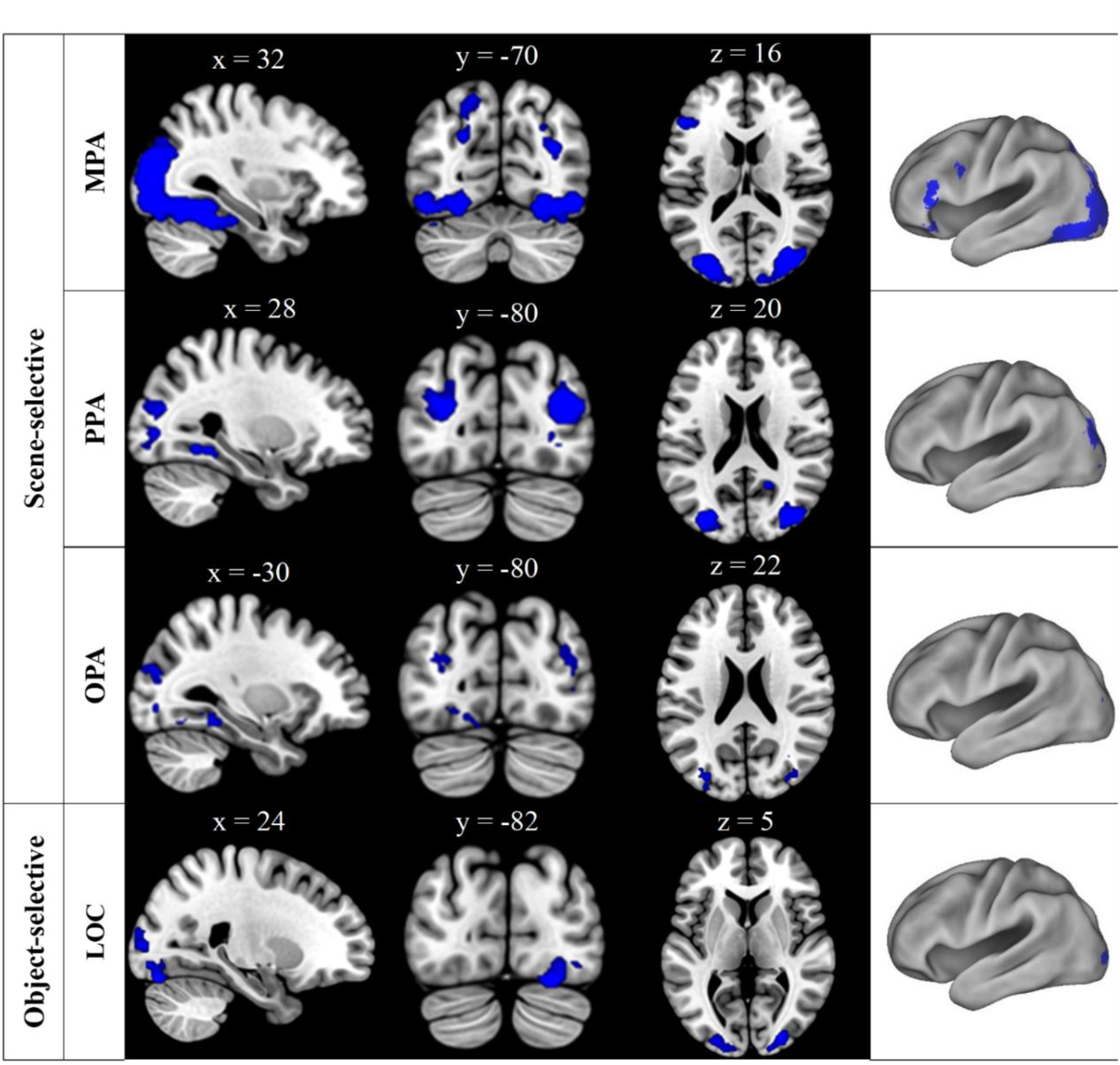
Functional connectivity effects at encoding for scene-selective seeds as a function of scene versus object trials that later received a correct source response, and for the object-selective LOC as a function of object versus scene trials that later received a correct source response. Clusters are displayed at height threshold: p < .001 (uncorrected) with cluster-level FWE correction (p < .05). **Alt text:** Brain images showing category-selective functional connectivity effects during encoding for scene-selective seeds and the object-selective LOC seed. Clusters are displayed on sagittal, coronal, axial, and surface-rendered views, mainly localized to posterior occipital and occipitotemporal regions.

**Table 3.** Coordinates of peak scene-selective connectivity effects at encoding.

| Region | MNI |  |  | Peak-z | K |
| --- | --- | --- | --- | --- | --- |
|  | x | y | z |  |  |
| Seed: MPA |  |  |  |  |  |
| Right middle occipital gyrus | 30 | -80 | 18 | 7.13 | 11024 |
| *Right fusiform gyrus\** | 30 | -54 | -12 | 6.96 |  |
| *Left middle occipital gyrus\** | -36 | -84 | 14 | 6.56 |  |
| *Left fusiform gyrus\** | -30 | -64 | -14 | 6.50 |  |
| *Right inferior occipital gyrus\** | 26 | -90 | -8 | 6.23 |  |
| Left inferior frontal gyrus | -52 | 32 | 12 | 5.38 | 500 |
| *Left inferior frontal gyrus* | -38 | 30 | -16 | 4.79 |  |
| Left inferior frontal sulcus | -46 | 8 | 30 | 3.94 | 164 |
| Seed: PPA |  |  |  |  |  |
| Right middle occipital gyrus | 38 | -80 | 18 | 6.18 | 728 |
| Left middle occipital gyrus | -28 | -82 | 18 | 5.82 | 616 |
| Left fusiform gyrus | -30 | -50 | -8 | 5.09 | 91 |
| Right precuneus | 16 | -56 | 16 | 5.05 | 109 |
| Right fusiform gyrus | 26 | -42 | -12 | 4.81 | 135 |
| Left inferior occipital gyrus | -18 | -90 | 0 | 4.39 | 122 |
| Seed: OPA |  |  |  |  |  |
| Left fusiform gyrus | -26 | -46 | -12 | 5.17 | 123 |
| Right fusiform gyrus | 26 | -56 | -8 | 4.97 | 178 |
| Left fusiform (posterior) gyrus | -24 | -76 | -10 | 4.80 | 168 |
| Right middle occipital gyrus | 36 | -74 | 26 | 4.52 | 155 |
| Left middle occipital gyrus | -34 | -82 | 20 | 4.45 | 110 |

Note: *Italicized* = subpeaks (which represent local maxima more than 4.0 mm apart from peak). Given the large extent of the MPA - occipito-temporal target cluster, anatomically distinct subpeaks (as defined by the AAL atlas) were identified within the cluster, and parameter estimates were extracted from these subpeaks (\*denotes subpeaks from which parameter estimates were extracted).

#### Object-selective effects

As shown in Figure 4 and detailed in Table 4, the LOC demonstrated enhanced connectivity with bilateral posterior occipital regions during subsequently remembered objects relative to scenes.

**Table 4.** Coordinates of object-selective connectivity effects at encoding.

| Region | MNI |  |  | Peak-z | K |
| --- | --- | --- | --- | --- | --- |
|  | x | y | z |  |  |
| Seed: LOC |  |  |  |  |  |
| Right lateral occipital gyrus | 24 | -82 | -18 | 5.33 | 687 |
| Right lingual gyrus | 14 | -94 | -6 | 4.92 |  |
| Right middle occipital gyrus | 28 | -90 | 8 | 4.81 |  |
| Left middle occipital gyrus | -26 | -96 | 2 | 4.48 | 264 |

Note: Italicized = subpeaks (which represent local maxima more than 4.0 mm apart from peak). K = cluster extent.

### Age differences in functional connectivity at encoding

#### Scene-selective effects

To examine whether scene-selective connectivity change at encoding varied as a function of target region and age group, mixed-design ANOVAs were conducted separately for each seed (see Table 5). For the MPA, there was a significant main effect of age group and a significant age group x target interaction. Follow-up independent-samples t-tests revealed that younger adults (M = .330, SD = .344) showed greater connectivity between the MPA and the left inferior frontal sulcus than older adults (M = .068, SD = .295), t(49) = 2.918, p_corrected =_ .030. Analyses of the PPA revealed a significant main effect of target region and a significant age group x target region interaction, but no main effect of age group.

**Table 5.** Results of mixed-design ANOVAs examining the effects of age group and target region on scene-selective functional connectivity change at encoding.

| Seed (# of targets) | F(df<sub>1</sub>, df<sub>2</sub>) | $\eta^2$ | p |
| --- | --- | --- | --- |
| **MPA (7)** |  |  |  |
| Target | 2.081(4.09, 200.18) | .041 | .083 |
| Age Group | 8.476(1, 49) | .147 | **.005** |
| Target x Age Group | 3.369(4.09, 200.18) | .064 | **.010** |
| **PPA (6)** |  |  |  |
| Target | 6.679(4.08, 200) | .120 | **<.001** |
| Age Group | .115(1, 49) | .002 | .736 |
| Target x Age Group | 2.654(4.08, 200) | .051 | **.033** |
| **OPA (5)** |  |  |  |
| Target | 1.261(4, 196) | .025 | .287 |
| Age Group | .235(1, 49) | .005 | .630 |
| Target x Age Group | 2.312(4, 196) | .045 | .059 |

Note: df = degrees of freedom, $\eta^2$ = partial eta squared. Significant effects ( $p < .05$ ) are in bold.

Independent samples t-test revealed that younger adults (M = .223, SD = .175) showed greater connectivity change between the PPA and left occipital pole than older adults (M = .048, SD = .273), t(49) = 2.683, p = .010, but this effect did not survive Holm-Bonferroni correction (p = .06). For the OPA, there were no significant main effects or interactions, indicating that encoding-related connectivity change of the OPA did not significantly differ across targets or between age groups. Figure 5 (panels A-C) illustrates the mean connectivity change between each scene-selective seed and its targets separately for younger and older adults.

**Figure 5.**
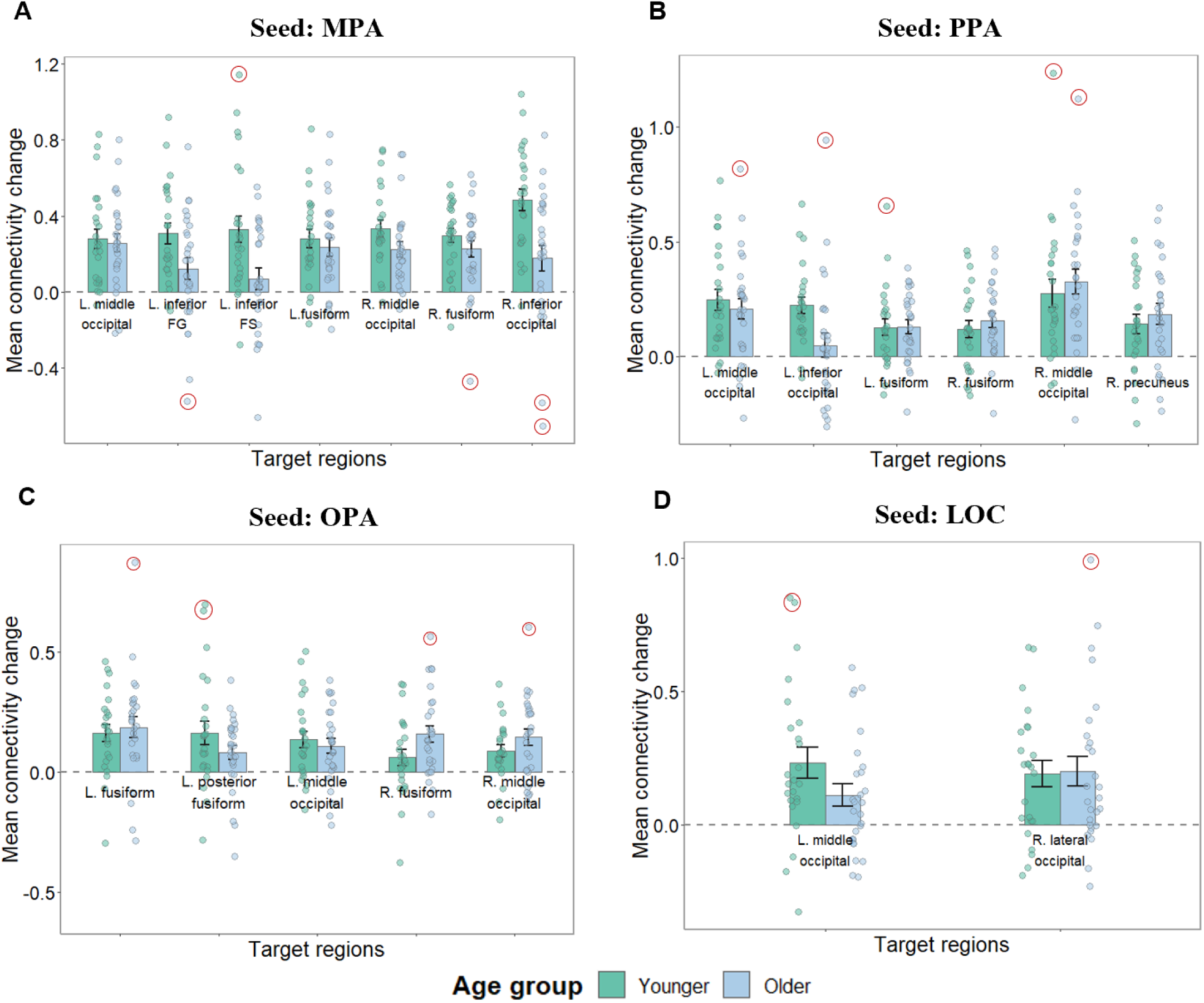
Mean connectivity change during scene- (panels A-C) and object- (panel D) selective encoding between each seed and its respective targets, shown separately for younger and older adults. Error bars represent standard errors of the mean. Outliers (+/- 2.5 SD from mean) are indicated by red circles. Repeating the analyses after winsorization of extreme values did not change the results. **Alt text:** Bar graphs showing mean category-selective connectivity change during encoding between scene-selective seeds and the object-selective LOC seed and their target regions in younger and older adults.

#### Object-selective effects

Analogous mixed-design ANOVAs were conducted to examine whether object-selective connectivity at encoding varied as a function of target region and age group (see Table 6). There were no significant main effects or interactions. Thus, LOC connectivity change during object encoding did not differ across target regions and was similar between younger and older adults. Figure 5 (panel D) illustrates the mean connectivity change between the LOC and its targets separately for younger and older adults.

**Table 6.**
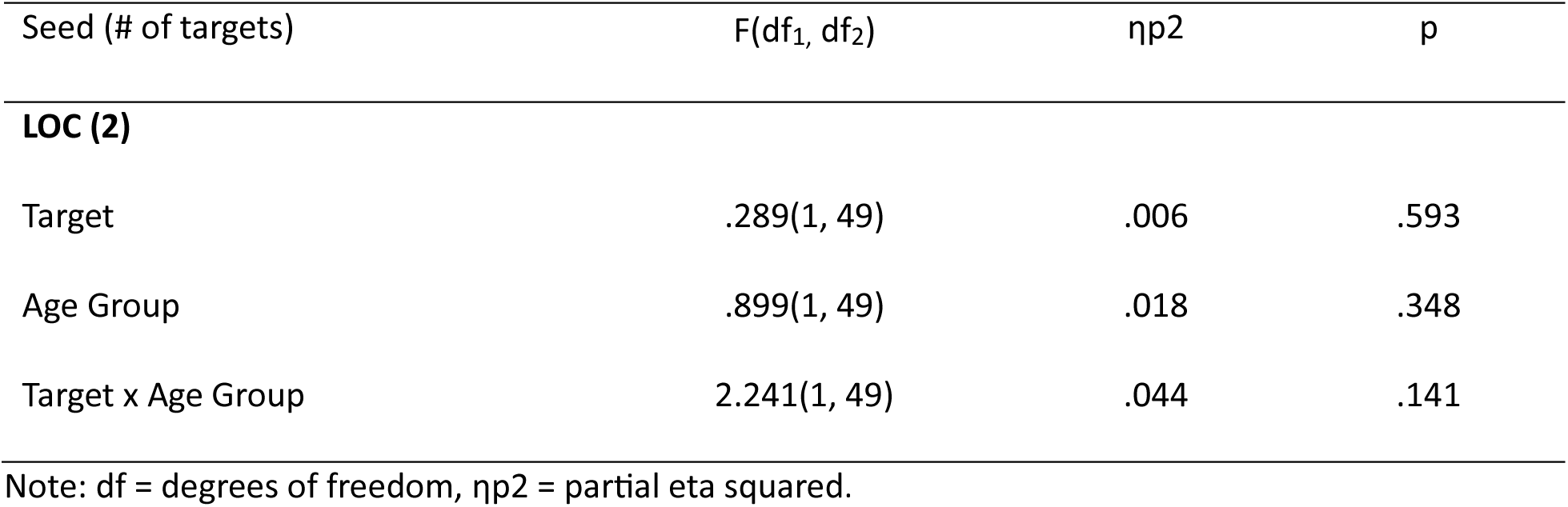
Results of mixed-design ANOVAs examining the effects of age group and target region on object-selective functional connectivity change at encoding.

### Overlap in functional connectivity effects at encoding

#### Scene-selective effects

To identify potential overlap in scene-selective connectivity effects at encoding, we inclusively masked the significant PPI main effects for each seed region that demonstrated a significant main effect (MPA, PPA, and OPA), thereby allowing the identification of convergent connectivity patterns. As shown in Figure 6, this analysis revealed overlapping connectivity effects across the scene-selective seeds. Overlapping clusters were identified in the right fusiform gyrus (73 voxels), right middle occipital gyrus (127 voxels), and left middle occipital gyrus (109 voxels).

**Figure 6.**
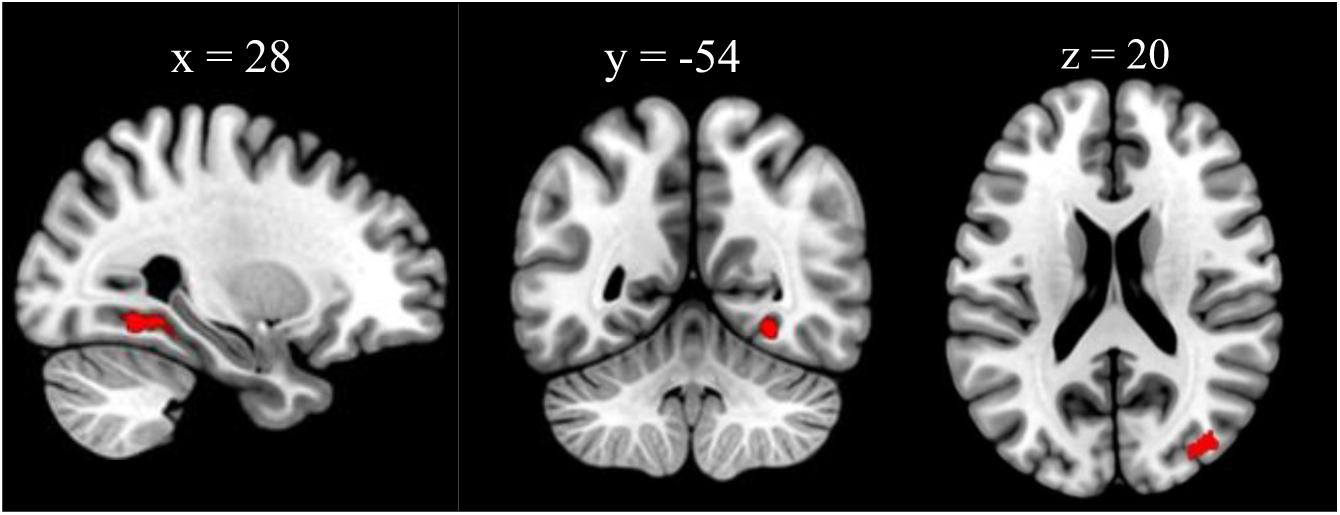
Inclusive masking of scene-selective PPI main effects at encoding revealed overlapping connectivity effects for the MPA, PPA, and OPA in the right fusiform gyrus and bilateral middle occipital gyri. Inclusive masking threshold: p < .001. **Alt text:** Three brain slices showing clusters in the right fusiform gyrus and bilateral middle occipital gyri on sagittal, coronal, and axial views.

#### Object-selective effects

Given that the LOC was the only seed showing significant connectivity change during object encoding, an overlap analysis analogous to that conducted for scene-selective seeds was not performed.

### Relationship between category-selective connectivity effects at encoding and subsequent memory performance

To investigate whether category-selective increases in functional connectivity during encoding were associated with subsequent memory performance, we conducted a series of multiple regression analyses using PPI-derived connectivity estimates extracted separately for each target region. Separate regression models were constructed predicting memory performance (source memory: pSR and item memory: Pr) using three predictors: age group, the PPI interaction parameter estimate, and their interaction. Given the high correlation between scene and object item memory (r = .910, p < .001), item memory scores were averaged across the two image categories. In cases where the age group x PPI interaction term did not reach statistical significance (p > .05), the term was removed from the model. Significant interaction effects were followed up with partial correlations computed separately for each age group while controlling for chronological age. All p-values were subjected to Holm-Bonferroni correction.

As detailed in Table 7 (see Supplementary Table 4 for a complete list including non-significant associations) regression analyses revealed that increased connectivity between scene-selective seeds and posterior cortical regions at encoding was associated with higher subsequent source memory performance, and that these associations were moderated by age group. As illustrated in Figure 7, follow-up partial correlations computed separately for each age group indicated that these interactions were driven by a positive association between connectivity change and memory performance in younger adults, with no significant association in older adults. However, in one case (OPA-right fusiform gyrus), the partial correlations did not reach significance in either age group [younger adults (r_partial_ = .301, p = .163); older adults (r_partial_ = -.185, p = .365)].

**Figure 7.**
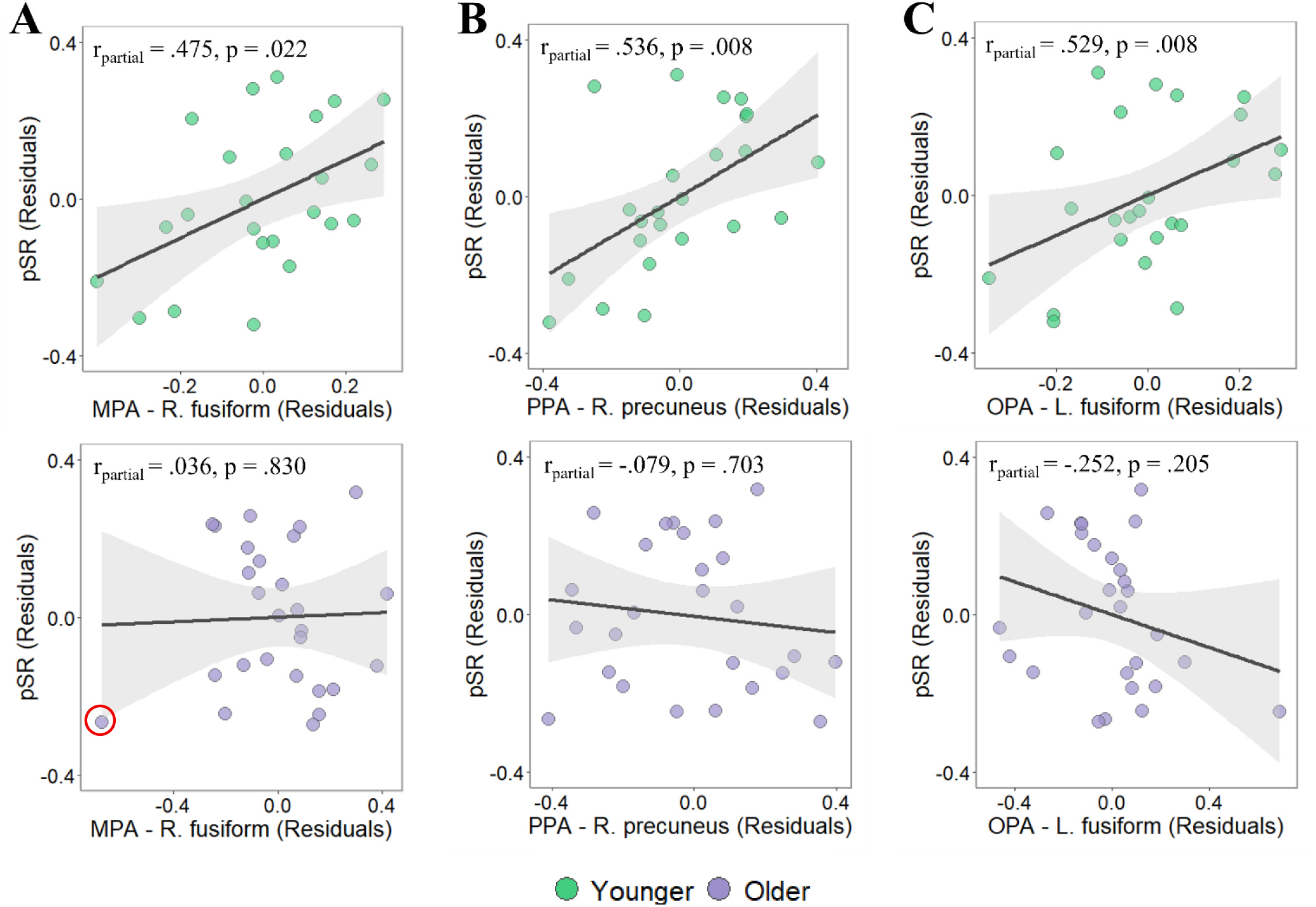
Scatterplots illustrating partial correlations between source memory performance (pSR) and scene-selective connectivity at encoding for (A) MPA - right fusiform gyrus, (B) PPA - right precuneus, and (C) OPA - left fusiform gyrus. Results are displayed separately for younger (top row) and older adults (bottom row), controlling for chronological age. Repeating the analyses after winsorization of extreme values (indicated by red circle) did not change the results The association between MPA-right fusiform connectivity and pSR did not survive Holm-Bonferroni correction. **Alt text:** Scatterplots showing relationships between source memory performance and scene-selective connectivity for younger and older adults. Younger adults showed positive associations between connectivity and source memory across all three seed-target pairs, whereas older adults showed non-significant associations. Regression lines are displayed for each scatterplot.

**Table 7.**
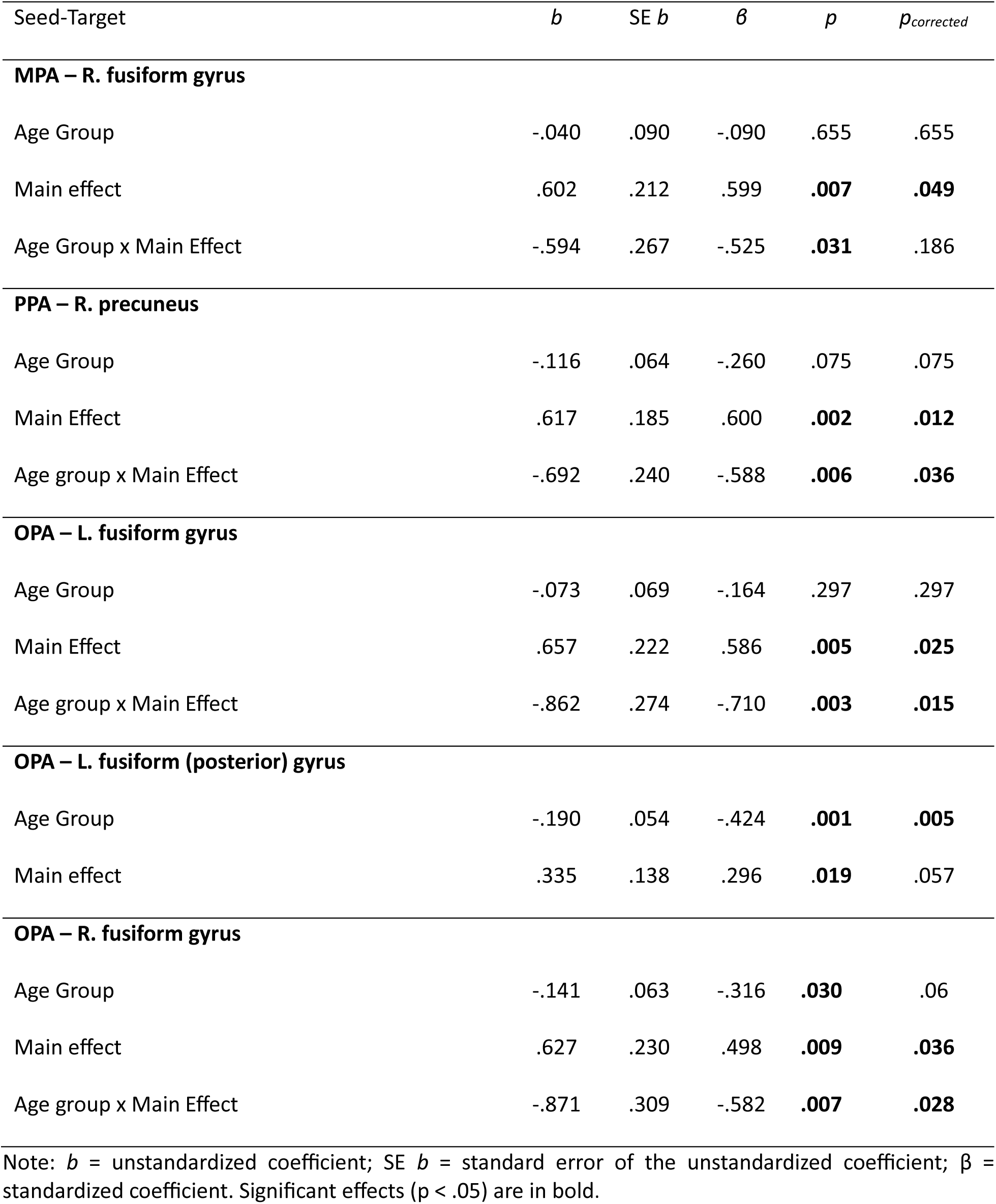
Results of the multiple regression analyses predicting source memory performance from age group and PPI main effects (seed-target connectivity change) during scene encoding.

| Seed-Target | *b* | SE *b* | $\beta$ | *p* | *p*<sub>corrected</sub> |
| --- | --- | --- | --- | --- | --- |
| **MPA – R. fusiform gyrus** |  |  |  |  |  |
| Age Group | -.040 | .090 | -.090 | .655 | .655 |
| Main effect | .602 | .212 | .599 | **.007** | **.049** |
| Age Group x Main Effect | -.594 | .267 | -.525 | **.031** | .186 |
| **PPA – R. precuneus** |  |  |  |  |  |
| Age Group | -.116 | .064 | -.260 | .075 | .075 |
| Main Effect | .617 | .185 | .600 | **.002** | **.012** |
| Age group x Main Effect | -.692 | .240 | -.588 | **.006** | **.036** |
| **OPA – L. fusiform gyrus** |  |  |  |  |  |
| Age Group | -.073 | .069 | -.164 | .297 | .297 |
| Main Effect | .657 | .222 | .586 | **.005** | **.025** |
| Age group x Main Effect | -.862 | .274 | -.710 | **.003** | **.015** |
| **OPA – L. fusiform (posterior) gyrus** |  |  |  |  |  |
| Age Group | -.190 | .054 | -.424 | **.001** | **.005** |
| Main effect | .335 | .138 | .296 | **.019** | .057 |
| **OPA – R. fusiform gyrus** |  |  |  |  |  |
| Age Group | -.141 | .063 | -.316 | **.030** | .06 |
| Main effect | .627 | .230 | .498 | **.009** | **.036** |
| Age group x Main Effect | -.871 | .309 | -.582 | **.007** | **.028** |

Note: *b* = unstandardized coefficient; SE *b* = standard error of the unstandardized coefficient; $\beta$ = standardized coefficient. Significant effects (*p* < .05) are in bold.

Only one seed-target pair also showed a significant association with item memory, namely, the PPA - right precuneus (β = .409, SE = .091, p = .003). No significant associations were observed between encoding-related LOC connectivity change and either pSR (see Supplementary Table 5) or Pr.

### Category-selective connectivity effects at retrieval

#### Scene-selective retrieval

As illustrated in Figure 8 and reported in Table 8, the scene-selective MPA exhibited increased connectivity with left middle and inferior frontal gyri, left medial superior frontal gyrus, and left intraparietal sulcus during successful scene retrieval. Similarly, the PPA demonstrated increased connectivity with left middle and inferior frontal gyri.

**Figure 8.**
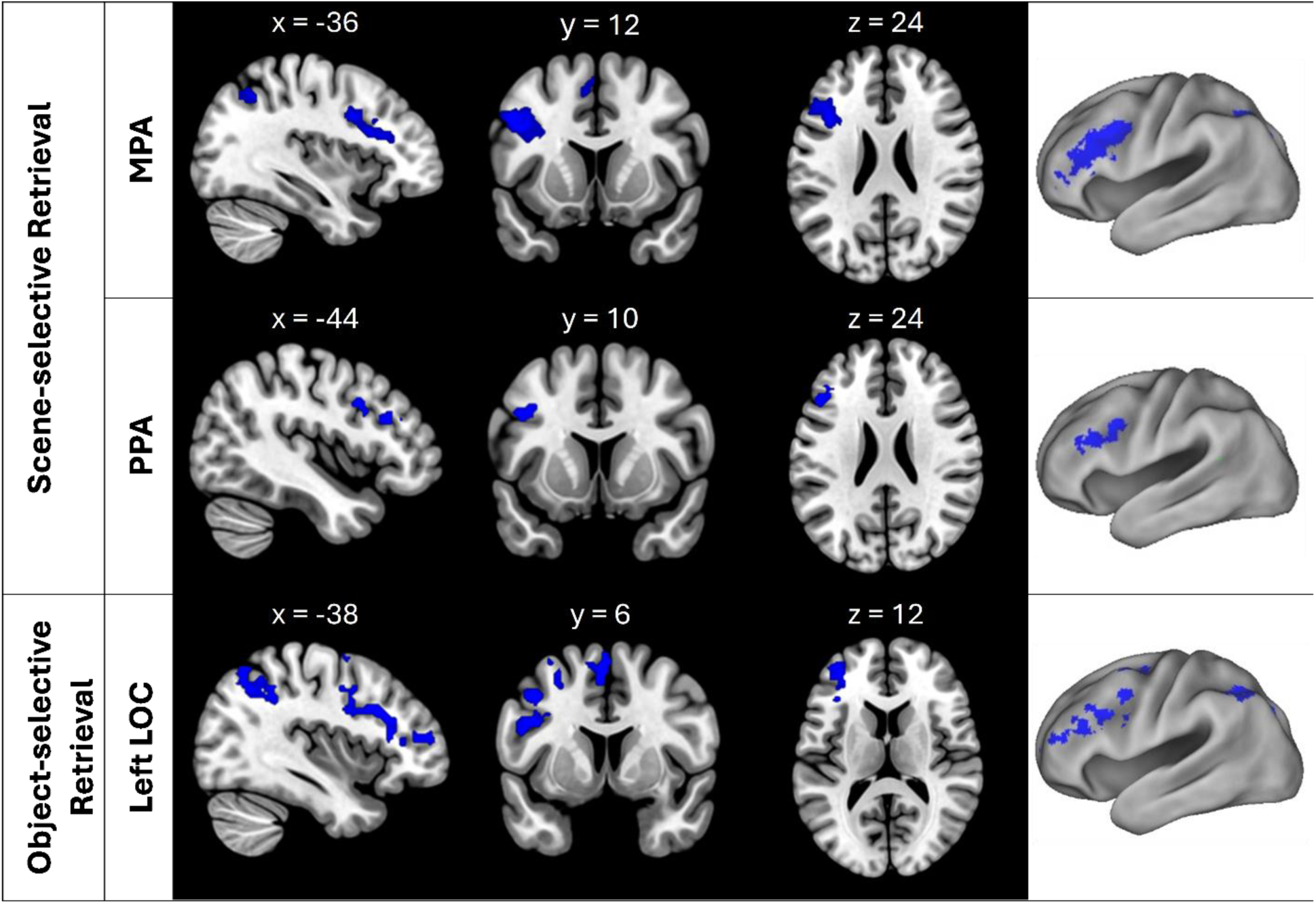
Functional connectivity change in scene-selective seeds as a function of scene versus object trials associated with a correct source memory judgment and in object-selective LOC as a function of object versus scene trials associated with a correct source memory judgment. Clusters are displayed at height threshold: p < .001 (uncorrected) with cluster-level FWE correction (p < .05). **Alt text:** Brain images showing category-selective functional connectivity effects during retrieval for scene-selective and object-selective seeds. Clusters are displayed on sagittal, coronal, axial, and surface-rendered views, localized to lateral prefrontal and parietal cortices.

**Table 8.** Coordinates of peak scene-selective connectivity effects at retrieval.

| Region | MNI |  |  | Peak-z | K |
| --- | --- | --- | --- | --- | --- |
|  | x | y | z |  |  |
| Seed: MPA |  |  |  |  |  |
| Left middle frontal gyrus | -46 | 12 | 34 | 5.55 | 1026 |
| *Left inferior frontal gyrus* | -42 | 32 | 14 | 4.93 |  |
| *Left inferior frontal gyrus* | -40 | 18 | 24 | 4.79 |  |
| Left intraparietal sulcus | -32 | -58 | 44 | 4.95 | 299 |
| Left medial superior frontal gyrus | -4 | 28 | 40 | 4.24 | 130 |
| *Left supplementary motor cortex* | -6 | 14 | 50 | 4.10 |  |
| Seed: PPA |  |  |  |  |  |
| Left middle frontal gyrus | -44 | 8 | 34 | 4.04 | 76 |
| Left inferior frontal gyrus | -48 | 22 | 22 | 3.99 | 83 |

Note: Italicized = subpeaks (which represent local maxima more than 4.0 mm apart from peak).

#### Object-selective retrieval

Analogous analyses investigating modulations in functional connectivity during object-selective retrieval revealed that the left LOC exhibited increased connectivity with left middle and inferior frontal gyri and left intraparietal sulcus (see Figure 8 and Table 9).

**Table 9.** Coordinates of peak object-selective connectivity effects at retrieval.

| Region | MNI |  |  | Peak-z | K |
| --- | --- | --- | --- | --- | --- |
|  | x | y | z |  |  |
| Seed: Left LOC |  |  |  |  |  |
| Left inferior frontal gyrus | -40 | 20 | 28 | 5.34 | 928 |
| *Left middle frontal gyrus* | -36 | 46 | 10 | 4.19 |  |
| Left intraparietal sulcus | -32 | -62 | 46 | 4.97 | 949 |
| *Left inferior parietal lobule* | -42 | -56 | 60 | 4.31 |  |
| Left supplementary motor area | -4 | 8 | 54 | 4.76 | 362 |
| Right cerebellum | 28 | -66 | -30 | 4.45 | 131 |
| Left precuneus | -12 | -70 | 38 | 4.41 | 148 |
| *Left precuneus* | -6 | -72 | 50 | 4.33 |  |
| Left middle frontal gyrus | -30 | 8 | 52 | 4.35 | 112 |

Note: Italicized = subpeaks (which represent local maxima more than 4.0 mm apart from peak).

### Age differences in functional connectivity effects at retrieval

#### Scene-selective effects

As for the analyses performed at encoding, mixed-design ANOVAs were conducted separately for each seed to examine whether connectivity at retrieval varied as a function of target region and age group (see Table 10). For the MPA, the ANOVA revealed a significant main effect of target, indicating that connectivity change differed across target regions, as well as a significant main effect of age group, reflecting greater overall connectivity change in older relative to younger adults. In contrast, analyses of PPA connectivity revealed no significant main effects or interactions, indicating that retrieval-related connectivity change with the PPA did not differ reliably across target regions or age groups. Figure 9 (panels A and B) illustrates the mean connectivity change between each scene-selective seed and its target regions separately for younger and older adults.

**Figure 9.**
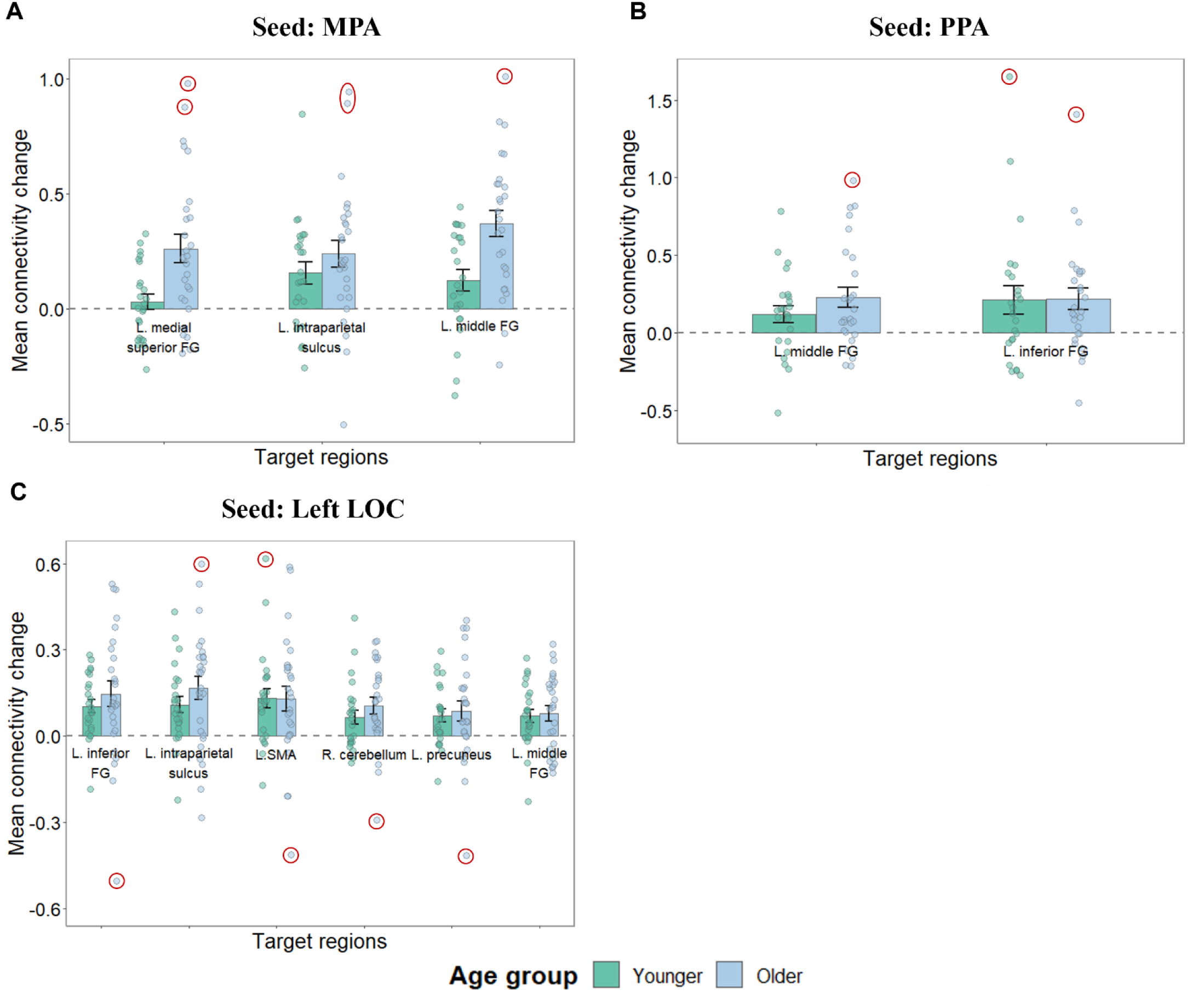
Mean connectivity change during scene retrieval (panels A and B) and object retrieval (panel C) between each seed and its target regions, shown separately for younger and older adults. Error bars represent standard error of the mean. Outliers (+/- 2.5 SD from the mean) are indicated by red circles. Repeating the analyses after winsorization of extreme values did not change the results. **Alt text:** Bar graphs showing mean category-selective connectivity change in scene- and object-selective ROIs during retrieval and their target regions in younger and older adults.

**Table 10.**
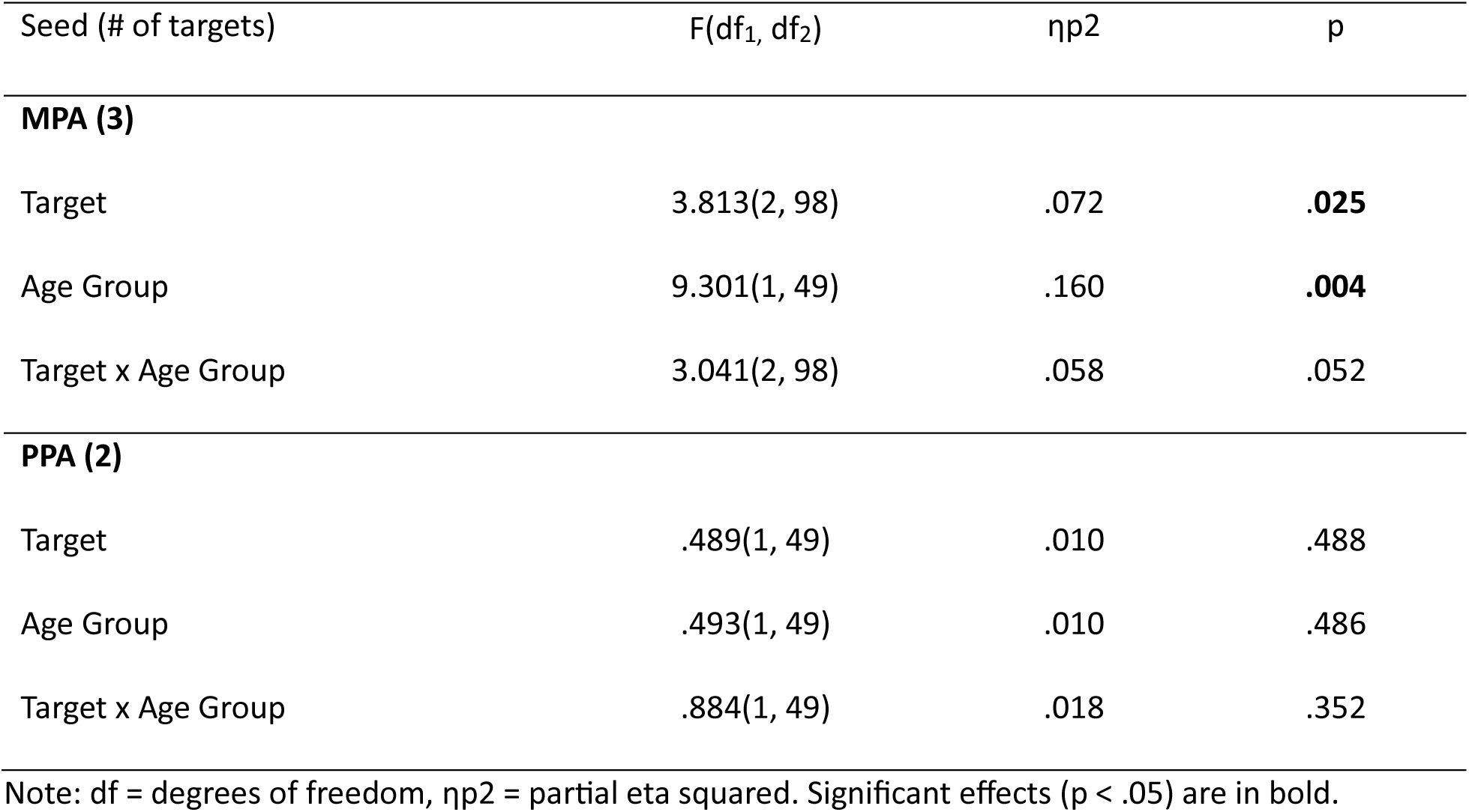
Results of mixed-design ANOVAs examining the effects of age group and target region on scene-selective functional connectivity change at retrieval.

#### Object-selective effects

An analogous mixed-design ANOVA revealed no significant main effects or interactions (see Table 11), indicating that left LOC connectivity change during retrieval did not vary reliably as a function of target region or age group. Figure 9 (panel C) illustrates the mean connectivity change between left LOC and its target regions separately for younger and older adults.

**Table 11.** Results of mixed-design ANOVAs examining the effects of age group and target region on object-selective functional connectivity change at retrieval.

| Seed (# of targets) | F(df<sub>1</sub>, df<sub>2</sub>) | $\eta^2$ | p |
| --- | --- | --- | --- |
| **Left LOC (6)** |  |  |  |
| Target | 2.387(3.88, 190.14) | .046 | .054 |
| Age Group | .760(1, 49) | .015 | .387 |
| Target x Age Group | .395(3.88, 190.14) | .008 | .807 |

Note: df = degrees of freedom, $\eta^2$ = partial eta squared.

### Overlap in functional connectivity effects at retrieval

#### Scene-selective effects

To identify regions of overlap in scene-selective connectivity during retrieval, we performed an inclusive masking analysis of the seed regions that demonstrated significant main effects in the initial retrieval-related connectivity analyses. Specifically, the PPI main effects of two scene-selective seeds (MPA and PPA) were inclusively masked. As illustrated in Figure 10, overlapping connectivity effects between the MPA and PPA were identified in the left middle frontal gyrus (75 voxels) and left inferior frontal gyrus (77 voxels).

**Figure 10.**
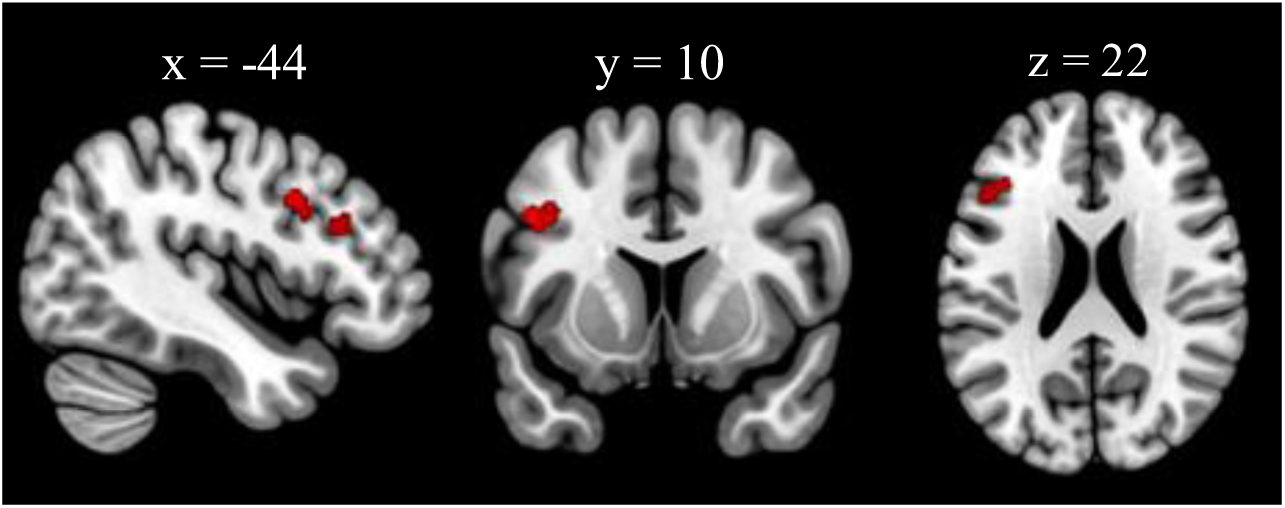
Inclusive masking of scene-selective PPI main effects at retrieval revealed overlapping connectivity effects for the MPA and PPA in the left lateral prefrontal cortex. Inclusive masking threshold: p < .001. **Alt text:** Three brain slices showing clusters in the left lateral prefrontal cortex on sagittal, coronal, and axial views.

#### Object-selective effects

An analogous inclusive masking analysis was not conducted for the left LOC given that it was the only object-selective seed examined.

### Relationship between category-selective connectivity effects at retrieval and memory performance

As is summarized in Table 12 (see Supplementary Table 6 for a complete list including non-significant associations), increased connectivity between the MPA and left middle frontal gyrus during scene-selective retrieval was positively associated with source memory performance, and this association was moderated by age group. As illustrated in Figure 11, follow-up partial correlations conducted separately for younger and older adults (controlling for chronological age) indicated that the association was significant in younger but not in older adults. No significant associations were observed between scene-selective connectivity at retrieval and item memory performance.

**Figure 11.**
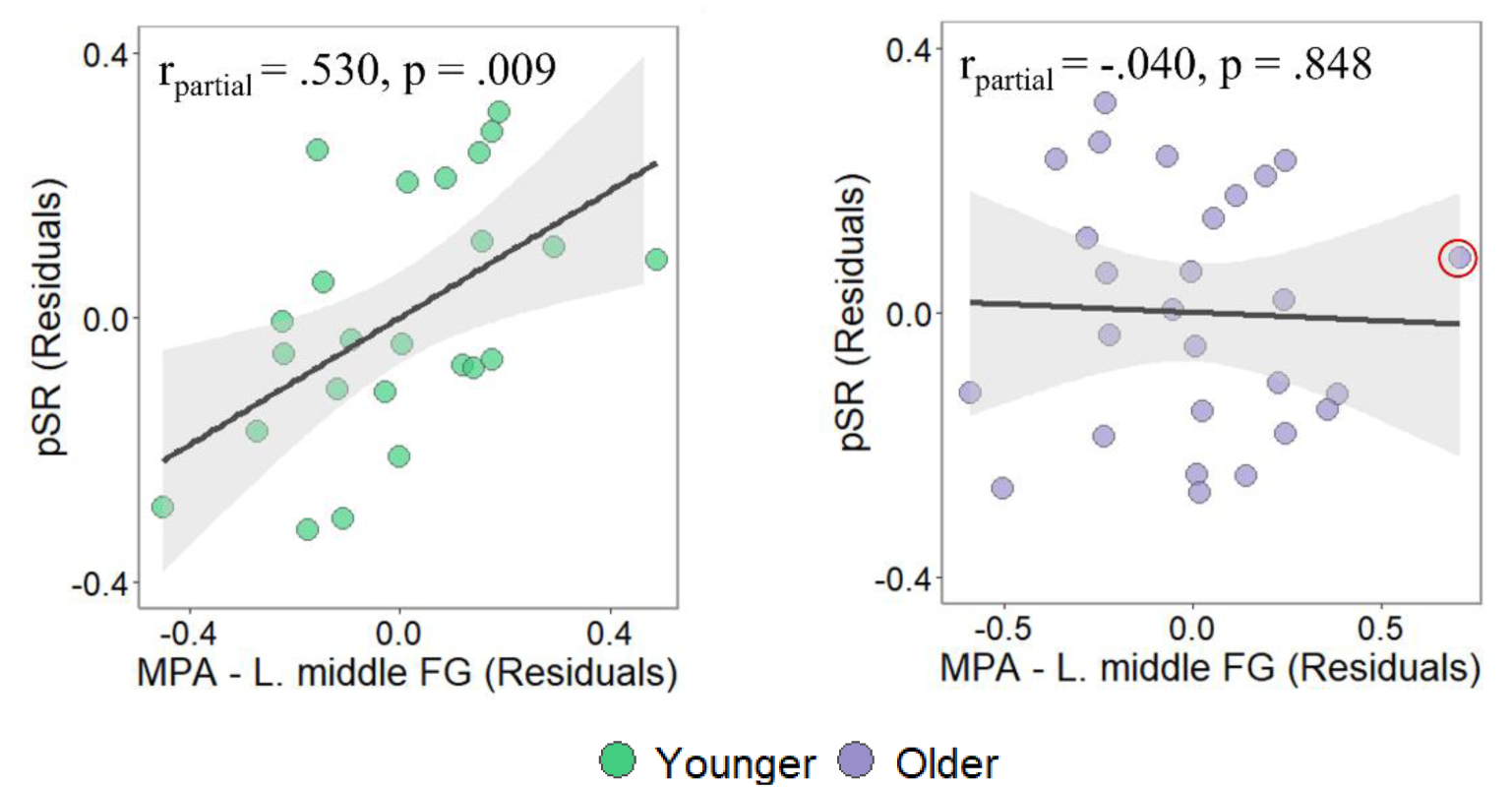
Scatterplots illustrating partial correlations between source memory and MPA –left middle frontal gyrus connectivity during scene retrieval. Repeating the analyses after winsorization of extreme values (indicated by red circles) did not change the results. **Alt text:** Scatterplots showing relationships between source memory performance and scene-selective connectivity for younger and older adults. Younger adults showed a positive association between connectivity and source memory, whereas older adults showed no significant association. Regression lines are displayed for each scatterplot.

**Table 12.**
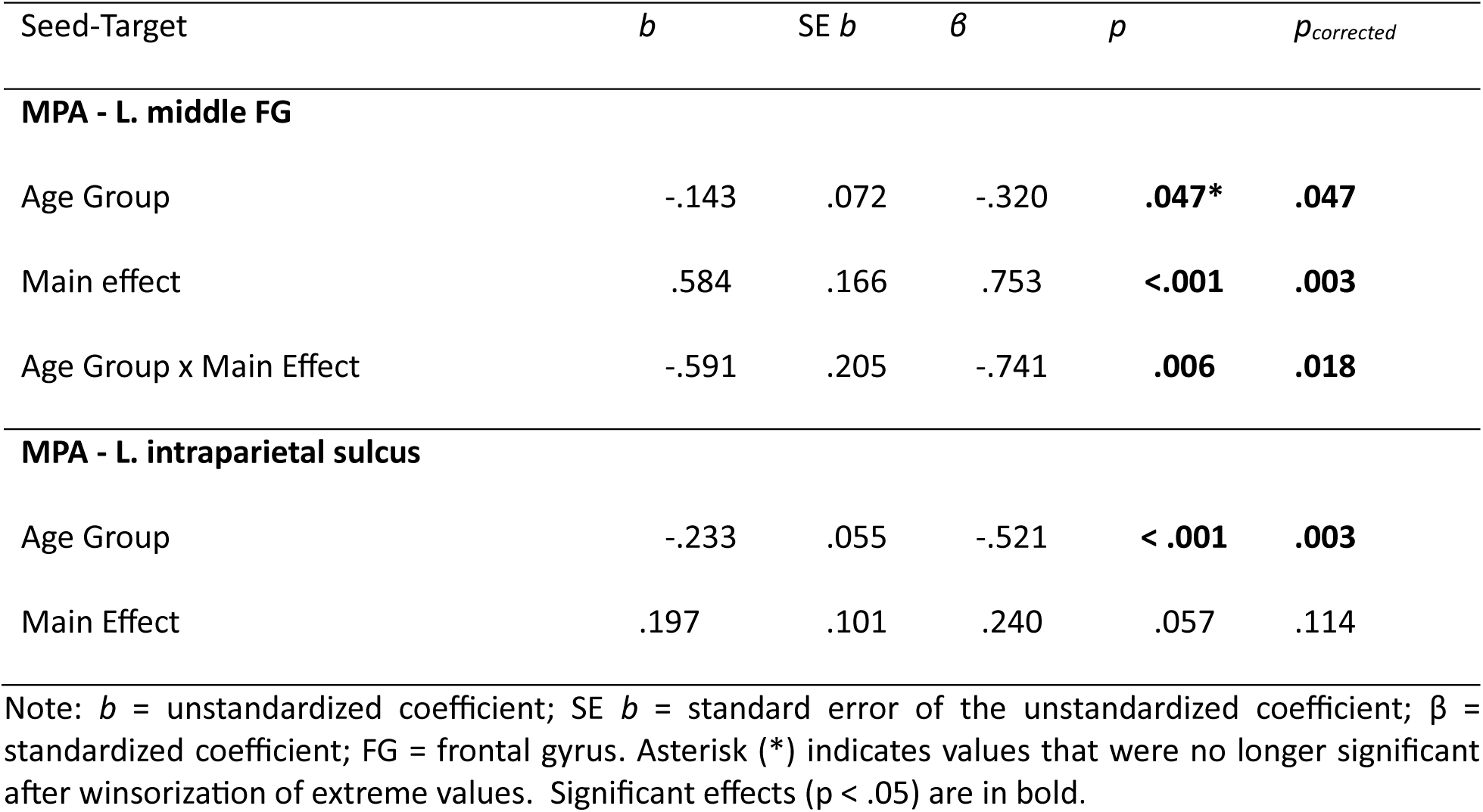
Results of the multiple regression analyses predicting source memory performance from age group and PPI main effects (seed-target connectivity change) during scene retrieval.

| Seed-Target | *b* | SE *b* | $\beta$ | *p* | *p*<sub>corrected</sub> |
| --- | --- | --- | --- | --- | --- |
| **MPA - L. middle FG** |  |  |  |  |  |
| Age Group | -.143 | .072 | -.320 | **.047\*** | **.047** |
| Main effect | .584 | .166 | .753 | **<.001** | **.003** |
| Age Group x Main Effect | -.591 | .205 | -.741 | **.006** | **.018** |
| **MPA - L. intraparietal sulcus** |  |  |  |  |  |
| Age Group | -.233 | .055 | -.521 | **< .001** | **.003** |
| Main Effect | .197 | .101 | .240 | .057 | .114 |

Note: *b* = unstandardized coefficient; SE *b* = standard error of the unstandardized coefficient; $\beta$ = standardized coefficient; FG = frontal gyrus. Asterisk (\*) indicates values that were no longer significant after winsorization of extreme values. Significant effects ( $p < .05$ ) are in bold.

As summarized in Table 13 (see Supplementary Table 7 for a complete list including non-significant associations), a significant main effect and an age group x connectivity interaction were observed for the association between pSR and left LOC-middle frontal gyrus connectivity. However, neither effect survived Holm-Bonferroni correction. No significant associations were observed between object-selective connectivity at retrieval and item memory performance.

**Table 13.**
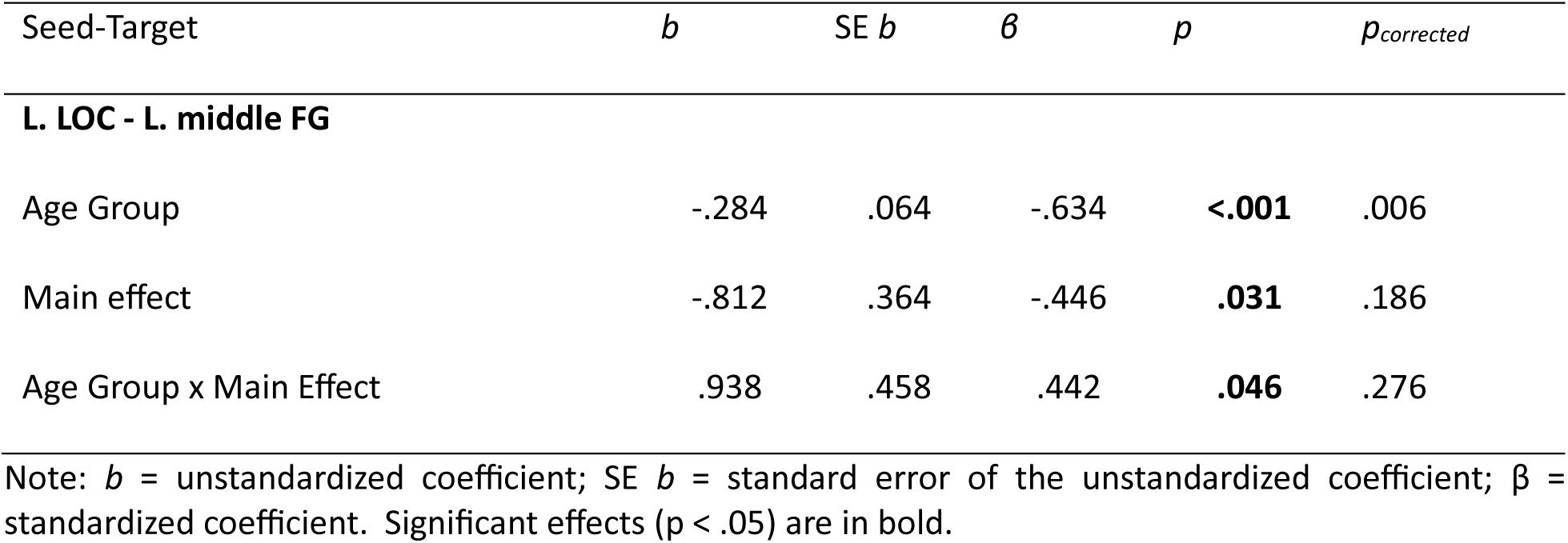
Results of the multiple regression analyses predicting source memory performance from age group and PPI main effects (seed-target connectivity change) during object retrieval.

## Discussion

In an extension of prior work that largely focused on activation-based measures of category selectivity, the present findings indicate that scene- and object-selective neural regions demonstrate partially converging patterns of connectivity during episodic encoding and retrieval. Scene-selective regions showed robust increases in connectivity with posterior occipital and occipitotemporal cortex during scene relative to object encoding. Similarly, during object relative to scene encoding the object selective LOC exhibited greater connectivity with posterior occipital and right occipitotemporal cortices. At retrieval, scene recall (relative to object recall) was characterized by increased connectivity between scene-selective regions and left lateral prefrontal and parietal cortices, and object recall was similarly associated with increased connectivity between the left LOC and left lateral prefrontal and parietal regions. Age differences in scene-selective connectivity strength were identified at both encoding and retrieval. Additionally, several scene-selective connectivity effects were associated with individual differences in source memory performance, and these associations varied as a function of age.

Prior to discussing the findings, we note an important consideration concerning the PPI contrasts employed in the present analyses. Encoding and retrieval connectivity effects were identified by contrasts between scene and object trials associated with accurate source memory judgments, rather than contrasts between trials associated with accurate versus unsuccessful source memory.

Consequently, the findings should be interpreted as reflecting category-selective modulation of connectivity associated with successful episodic memory, rather than neural correlates of successful versus unsuccessful retrieval.

### Encoding

Both scene- and object-selective regions exhibited enhanced connectivity with posterior visual regions during the encoding of their preferred content. In addition, and consistent with the resting state findings reported by Stevens et al., (2015), we identified increased inter-regional connectivity between different scene-selective regions during scene encoding, most notably between the MPA and PPA. During scene encoding the MPA also exhibited increased connectivity with the left lateral prefrontal cortex (latPFC), including the inferior and middle frontal gyri. Prior research has consistently implicated these and adjacent regions in processes that support associative encoding (e.g., Buckner et al., 1999; Murray & Ranganath, 2007; Park & Rugg, 2008; Blumenfeld et al., 2011; Park & Rugg, 2011; Wong et al., 2013).

One interpretation of the present findings is that scene encoding places relatively greater demands on associative encoding processes than does object encoding. Although both scene and object trials required the formation of word-image associations, scenes comprise multiple spatial and contextual elements that must be integrated into a unified scene representation.

Why was connectivity with left latPFC observed for the MPA but not for other scene-selective regions? One possibility arises from the findings from resting-state fMRI studies which suggest that scene-selective cortex is organized into two distinct large-scale networks (Baldassano et al., 2016; Watson & Andrews, 2024). Notably, the MPA is proposed to belong to a network that supports the integration of incoming scene information with stored spatial and contextual knowledge. By this account, the present findings of dissociable connectivity between scene regions and latPFC may, in part, reflect this broader functional organization of scene-selective cortex.

Relative to older adults, younger adults showed greater scene-selective connectivity effects between the MPA and left inferior frontal sulcus. In contrast, object-selective connectivity at encoding was seemingly age-invariant. Prior studies have reported age-related differences in encoding-related functional connectivity (Grady et al., 2003; Dennis et al., 2008; Cansino et al., 2017; Ankudowich et al., 2018). However, many of these studies primarily assessed connectivity between the hippocampus and other medial temporal regions with prefrontal and/or posterior cortical regions.

One interpretation for why age differences were found for scene but not object connectivity at encoding is that the neural correlates of scene processing may be more sensitive to aging effects than the correlates of object processing. Consistent with this idea, prior work suggests that older adults process complex visual inputs, such as natural scenes, that contain multiple spatially distributed elements differently than younger adults (Agnew & Pilz, 2017; Rycroft et al., 2018; Bouhassoun et al., 2022). Evidence from studies of age-related neural dedifferentiation also suggest that the neural mechanisms underlying the encoding of different image categories may vary in their sensitivity to aging. Specifically, category-level dedifferentiation is largely restricted to scene-selective regions (Koen et al., 2019, 2020; Srokova et al., 2020; 2024). While the present findings concern functional connectivity rather than neural specificity, this literature is broadly consistent with the idea that the neural correlates of scene encoding may be particularly sensitive to age differences.

Age-related differences were also evident in the relationships between encoding-related connectivity and subsequent memory performance. Across several scene-selective seed-target pairs, greater scene-related connectivity change was associated with higher subsequent source memory performance in younger adults, whereas no reliable associations were observed in older adults.

Importantly, the seed-target pairs that showed age differences in connectivity strength were distinct from those that were associated with subsequent memory performance in young adults. Thus, these age-moderated connectivity-behavior associations cannot be explained simply by age differences in connectivity strength.

#### Retrieval

Successful scene retrieval was associated with increased connectivity between the MPA and the left latPFC, including middle and inferior frontal gyri, as well as with left intraparietal sulcus. The PPA exhibited a similar, albeit less extensive, pattern of connectivity with left middle and inferior frontal gyri, and at a less stringent threshold (p< .005), with the left lateral parietal cortex also. Of importance, MPA and PPA connectivity converged on overlapping left latPFC regions during scene retrieval. Object retrieval was similarly associated with increased connectivity between the left LOC and left latPFC and intraparietal sulcus.

Both lateral prefrontal and parietal cortices comprise multiple architectonically distinct regions that differ in their connectivity profiles and functional characteristics (Johansen-Berg et al., 2004; Petrides, 2005; Caspers et al., 2006; Wang et al., 2017; Passingham & Wise, 2012). The latPFC has been implicated in several retrieval-related processes, including the selection of task-relevant mnemonic representations, resolution of competition among retrieved representations, and the monitoring and evaluation of retrieved information (see Rugg, 2024, for a review; see also Wagner et al., 2005; Badre et al., 2005; Badre & Wagner, 2007; Chadick & Gazzaley, 2011; Barredo et al., 2015; Hutchinson et al., 2014).

Increased connectivity between latPFC and other cortical regions during episodic retrieval has been observed in prior studies. For example, Kostopoulos and Petrides (2016) reported increased connectivity during episodic retrieval between the ventrolateral prefrontal cortex and superior temporal gyrus and sulcus as well as the inferior parietal lobule although these effects were right-lateralized.

Similarly, King et al. (2015) reported increased connectivity between regions held to belong to a “core recollection network” and left lateral prefrontal and parietal cortices during successful recollection, findings which they interpreted as reflecting the engagement of control processes supporting post-retrieval operations.

One interpretation of the retrieval-related connectivity effects identified between category-selective regions and left lateral prefrontal and parietal cortices in the present study is that the effects reflect engagement of control processes that support the retrieval of category-selective information. An important caveat, however, is that functional connectivity analyses do not provide information about the directionality of information flow. Consequently, the present findings are equally compatible with the engagement of top-down control processes (as we suggest above), bottom-up signaling from content-selective regions, or bidirectional effects. Nonetheless, the findings highlight the importance of connectivity-based analyses for characterizing the contribution of regions implicated in control processes to episodic retrieval beyond what can be inferred from regional activation alone.

As in the case of the encoding data, age differences were also evident in retrieval-related scene-selective connectivity. In striking contrast to the age effects identified at encoding, however, the MPA demonstrated greater connectivity effects with left lateral prefrontal regions in the older than in the younger adults. In contrast, object-selective connectivity was age-invariant, analogous to the pattern observed at encoding. Prior studies investigating age differences in task-related connectivity have employed a range of analytical approaches, seed regions, and task contrasts and focused mainly on connectivity involving the medial temporal lobe and prefrontal cortex (e.g. Grady et al., 2003; Daselaar et al., 2006; Dennis et al., 2008; Dew et al., 2012; Monge et al., 2018). Across these studies, one consistent finding is that older relative to younger age is associated with increased connectivity between posterior medial temporal lobe and prefrontal regions.

Prior work has suggested that increased connectivity between medial temporal or posterior cortical regions and prefrontal cortex in older adults may reflect compensatory recruitment of executive control processes to mitigate age-related declines in posterior regions supporting memory performance (Dennis et al., 2008; Dennis & Peterson, 2012), although these studies primarily examined connectivity at encoding. The present findings do not permit a straightforward interpretation of the increased MPA-prefrontal connectivity in older adults as being compensatory. If this connectivity pattern reflected a compensatory mechanism, arguably these connectivity metrics should have demonstrated a positive association with memory performance in older adults (Cabeza et al., 2018). However, we found no evidence for such an association. In contrast, among younger adults, increased connectivity between the MPA and left middle frontal gyrus was positively associated with source memory accuracy.

Indeed, analogous to the findings from the encoding phase, associations between retrieval-related connectivity change and memory performance were confined to scene trials and younger adults. The reasons for this pattern of findings are currently unclear, but we suspect that part of the explanation might lie with the unique attentional and mnemonic demands imposed by scene relative to object images, demands that are met less well by older adults (see Encoding subsection above).

### Limitations

Several limitations should be considered when interpreting the present findings. First, given the cross-sectional design of the study, age-related effects cannot be disentangled from potential cohort and selection effects (e.g. Rugg, 2016). Second, functional connectivity was assessed with PPI analyses, which capture task-dependent changes in connectivity but do not provide information about the directionality or causal influence between regions. Future work employing effective connectivity approaches may help clarify the directionality underlying these results. Finally, the relatively modest sample size may have limited sensitivity to detect additional age-related differences in category-selective connectivity. Replication in larger samples will be important for establishing the generalizability of the observed connectivity effects and connectivity-behavior relationships.

### Conclusions

The findings from the present study suggest that category-selective cortical regions exhibit partially convergent patterns of functional connectivity during episodic encoding and retrieval. These findings extend prior work that has largely examined category-selective regional activation effects.

During encoding, scene- and object-selective regions demonstrated increased connectivity with posterior occipital regions and during retrieval, with left lateral frontal and parietal regions. Age-related differences were observed for scene-selective connectivity at both encoding and retrieval. In addition, age-related differences were observed in the relationship between scene-selective connectivity and memory performance, such that greater connectivity was associated with higher source memory performance in younger adults only. Together, these findings highlight the value of task-based functional connectivity approaches for characterizing the neural mechanisms underlying category-selective episodic encoding and retrieval.

## Acknowledgments

We gratefully acknowledge the experimental volunteers who contributed their time to this study.

