## Supplementary Materials for "Category-selective functional connectivity during episodic encoding and retrieval in younger and older adults"

**Supplementary Table 1.** Coordinates of material-selective univariate encoding effects.

| Contrast | Region | MNI | | | Peak-z | K |
|---|---|---|---|---|---|---|
| | | x | y | z | | |
| **Scene-selective encoding** | Right fusiform gyrus | 28 | -46 | -8 | Inf | 25251 |
| | *Left fusiform/lingual gyrus* | *-26* | *-52* | *-8* | *Inf* | |
| | *Right precuneus* | *16* | *-56* | *16* | *Inf* | |
| | *Left precuneus* | *-18* | *-62* | *16* | *Inf* | |
| | *Right middle occipital gyrus* | *38* | *-80* | *22* | *Inf* | |
| | *Left calcarine gyrus* | *-6* | *-92* | *-2* | *Inf* | |
| | *Right lingual gyrus* | *6* | *-82* | *-4* | *Inf* | |
| | *Left middle occipital gyrus* | *-34* | *-86* | *16* | *Inf* | |
| | Left cerebellum | -14 | -46 | -50 | Inf | 979 |
| | *Right cerebellum* | *12* | *-48* | *-52* | *Inf* | |
| | Left subgenual cortex | 0 | 16 | -6 | 7.65 | 697 |
| | *Right orbitofrontal gyrus* | *4* | *44* | *-14* | *5.81* | |
| | Left superior temporal sulcus | -54 | -6 | -18 | 5.67 | 363 |
| | Right superior temporal sulcus | 56 | -4 | -18 | 5.61 | 244 |
| | Left middle frontal gyrus | -22 | 22 | 44 | 5.03 | 87 |
| | Right middle frontal gyrus | 26 | 20 | 42 | 4.77 | 362 |
| **Object-selective encoding** | Right inferior occipital gyrus | 52 | -66 | -12 | Inf | 2206 |
| | *Right fusiform gyrus* | *44* | *-46* | *-16* | *7.20* | |
| | Left inferior occipital gyrus | -48 | -70 | -6 | Inf | 2385 |

| | | | | | | |
|---|---|---|---|---|---|---|
| | *Left fusiform gyrus* | *-44* | *-50* | *-20* | *Inf* | |
| | Left supramarginal gyrus | -56 | -32 | 38 | 7.37 | 2274 |
| | Left inferior frontal gyrus | -38 | 34 | 12 | 6.41 | 995 |
| | *Left inferior frontal gyrus* | *-44* | *36* | *20* | *5.49* | |
| | Left precentral gyrus | -50 | 4 | 24 | 6.33 | 472 |
| | Right postcentral gyrus | 30 | -46 | 74 | 5.65 | 1451 |
| | *Right supramarginal gyrus* | *62* | *-28* | *44* | *5.33* | |
| | Right inferior frontal gyrus | 46 | 38 | 10 | 5.65 | 374 |
| | Left inferior frontal gyrus | -24 | 30 | -16 | 5.49 | 166 |
| | Right inferior frontal gyrus | 22 | 30 | -18 | 5.54 | 96 |
| | Left posterior insula | -40 | -6 | 2 | 5.28 | 113 |
| | Right middle temporal gyrus | 58 | -36 | 4 | 4.72 | 89 |
| | Right cerebellum | 20 | -84 | -44 | 4.16 | 108 |

Note: Italicized = subpeaks (which represent local maxima more than 4.0 mm apart from peak). K = cluster extent.

**Supplementary Table 2.** Coordinates of material-selective univariate retrieval effects.

| Contrast | Region | MNI | | | Peak-z | K |
|---|---|---|---|---|---|---|
| | | x | y | z | | |
| **Scene-selective retrieval** | Left precuneus | -12 | -58 | 12 | Inf | 1905 |
| | *Right precuneus* | *18* | *-58* | *20* | *Inf* | |
| | Right fusiform | 30 | -36 | -14 | Inf | 763 |
| | Left fusiform | -32 | -42 | -12 | Inf | 719 |
| | Left middle occipital gyrus | -36 | -84 | 30 | 5.98 | 475 |
| | Right middle occipital gyrus | 44 | -70 | 24 | 5.10 | 315 |
| | Left middle orbitofrontal gyrus | -12 | 44 | -6 | 4.46 | 263 |
| | Right superior medial frontal gyrus | 14 | 52 | 4 | 4.05 | |
| | Right orbitofrontal gyrus | 2 | 34 | -16 | 3.87 | |
| | Left middle temporal gyrus | -52 | -12 | -16 | 4.38 | 114 |
| **Object-selective retrieval** | Left inferior temporal gyrus | -44 | -58 | -6 | 3.87 | 102 |

Note: Italicized = subpeaks (which represent local maxima more than 4.0 mm apart from peak). K = cluster extent.

**Supplementary Table 3.** Coordinates of scene-selective connectivity effects at encoding.

| Region | MNI | | | Peak-z | K |
|---|---|---|---|---|---|
| | x | y | z | | |
| **Seed: MPA** | | | | | |
| Right middle occipital gyrus | 30 | -80 | 18 | 7.13 | 11024 |
| *Right fusiform gyrus** | *30* | *-54* | *-12* | *6.96* | |
| *Left middle occipital gyrus** | *-36* | *-84* | *14* | *6.56* | |
| *Left fusiform gyrus** | *-30* | *-64* | *-14* | *6.50* | |
| *Right inferior occipital gyrus** | *26* | *-90* | *-8* | *6.23* | |
| Left inferior frontal | -52 | 32 | 12 | 5.38 | 500 |
| *Left inferior frontal gyrus* | *-38* | *30* | *-16* | *4.79* | |
| Left inferior frontal sulcus | -46 | 8 | 30 | 3.94 | 164 |
| **Seed: PPA** | | | | | |
| Right middle occipital gyrus | 38 | -80 | 18 | 6.18 | 728 |
| *Right inferior occipital gyrus* | *26* | *-84* | *2* | *4.84* | |
| *Right inferior occipital gyrus* | *30* | *-88* | *-10* | *3.41* | |
| Left middle occipital gyrus | -28 | -82 | 18 | 5.82 | 616 |
| *Left middle occipital gyrus* | *-20* | *-96* | *16* | *3.62* | |
| Left fusiform gyrus | -30 | -50 | -8 | 5.09 | 91 |
| Right precuneus | 16 | -56 | 16 | 5.05 | 109 |
| Right fusiform | 26 | -42 | -12 | 4.81 | 135 |
| Left inferior occipital gyrus | -18 | -90 | 0 | 4.39 | 122 |
| **Seed: OPA** | | | | | |
| Left fusiform gyrus | -26 | -46 | -12 | 5.17 | 123 |

| | | | | | |
|---|---|---|---|---|---|
| Right fusiform/lingual gyrus | 26 | -56 | -8 | 4.97 | 178 |
| *Right fusiform gyrus* | *28* | *-28* | *-22* | *4.56* | |
| Left fusiform (posterior) gyrus | -24 | -76 | -10 | 4.80 | 168 |
| *Left lingual gyrus* | *-14* | *-84* | *-14* | *3.88* | |
| Right middle occipital gyrus | 36 | -74 | 26 | 4.52 | 155 |
| *Right middle occipital gyrus* | *38* | *-78* | *4* | *3.92* | |
| Left middle occipital gyrus | -34 | -82 | 20 | 4.45 | 110 |

Note: Italicized = subpeaks (which represent local maxima more than 4.0 mm apart from peak). Given the large extent of the MPA - occipito-temporal target cluster, anatomically distinct subpeaks (as defined by the AAL atlas) were identified within the cluster, and parameter estimates were extracted from these subpeaks (*denotes subpeaks from which parameter estimates were extracted).

**Supplementary Table 4.** Results of the multiple regression analyses predicting source memory performance from age group and PPI main effects (seed-target connectivity change) during scene encoding.

| Seed - Target | *b* | SE *b* | *β* | *p* | $p_{corrected}$ |
|---|---|---|---|---|---|
| **MPA - L. middle occipital gyrus** | | | | | |
| Age Group | -.214 | .056 | -.477 | **<.001** | **.007** |
| Main effect | .144 | .116 | .155 | .218 | .654 |
| **MPA - L. fusiform gyrus** | | | | | |
| Age Group | -2.07 | .055 | -.462 | **<.001** | **.007** |
| Main effect | .230 | .115 | .245 | .050 | .300 |
| **MPA - L. inferior FG** | | | | | |
| Age Group | -.210 | .059 | -.469 | **<.001** | **.007** |
| Main effect | .038 | .101 | .050 | .710 | 1.000 |
| **MPA - L. inferior FS** | | | | | |
| Age Group | -.210 | .061 | -.470 | **.001** | **.007** |
| Main effect | .025 | .090 | .038 | .783 | 1.000 |
| **MPA - R. middle occipital gyrus** | | | | | |
| Age Group | -.194 | .057 | -.434 | **.001** | **.007** |
| Main effect | .210 | .134 | .199 | .124 | .496 |
| **MPA - R. fusiform gyrus** | | | | | |
| Age Group | -.040 | .090 | -.090 | .655 | .655 |
| Main effect | .602 | .212 | .599 | **.007** | **.049** |
| Age Group x Main Effect | -.594 | .267 | -.525 | **.031** | .186 |
| **MPA – R. inferior occipital gyrus** | | | | | |
| Age Group | -.164 | .061 | -.367 | **.009** | **.018** |

| | | | | | |
|---|---|---|---|---|---|
| Main effect | .171 | .087 | .267 | .055 | .300 |
| **PPA – L. middle occipital gyrus** | | | | | |
| Age Group | -.214 | .057 | -.478 | **< .001** | **.006** |
| Main effect | .074 | .124 | .076 | .552 | 1.000 |
| **PPA – L. fusiform gyrus** | | | | | |
| Age Group | -.217 | .056 | -.485 | **< .001** | **.006** |
| Main effect | .052 | .354 | -.031 | .878 | 1.000 |
| **PPA – L. inferior occipital gyrus** | | | | | |
| Age Group | -.193 | .60 | -.430 | **.002** | **.006** |
| Main effect | .139 | .122 | .152 | .261 | 1.000 |
| **PPA – R. middle occipital gyrus** | | | | | |
| Age Group | -.220 | .057 | -.491 | **< .001** | **.006** |
| Main effect | .058 | .100 | .073 | .565 | 1.000 |
| **PPA – R. precuneus** | | | | | |
| Age Group | -.116 | .064 | -.260 | .075 | .075 |
| Main Effect | .617 | .185 | .600 | **.002** | **.012** |
| Age group x Main Effect | -.692 | .240 | -.588 | **.006** | **.036** |
| **PPA – R. fusiform gyrus** | | | | | |
| Age Group | -.229 | .055 | -.512 | **< .001** | **.006** |
| Main effect | .310 | .163 | .233 | .064 | .320 |
| **OPA – L. fusiform gyrus** | | | | | |
| Age Group | -.073 | .069 | -.164 | .297 | .297 |
| Main Effect | .657 | .222 | .586 | **.005** | **.025** |
| Age group x Main Effect | -.862 | .274 | -.710 | **.003** | **.015** |

| | | | | | |
|---|---|---|---|---|---|
| **OPA – L. fusiform (posterior) gyrus** | | | | | |
| Age Group | -.190 | .054 | -.424 | **.001** | **.005** |
| Main effect | .335 | .138 | .296 | **.019** | .057 |
| **OPA – L. middle occipital gyrus** | | | | | |
| Age Group | -.210 | .055 | -.469 | **<.001** | **.005** |
| Main effect | -.267 | .170 | .194 | .124 | .248 |
| **OPA – R. fusiform gyrus** | | | | | |
| Age Group | -.141 | .063 | -.316 | **.030** | .060 |
| Main effect | .627 | .230 | .498 | **.009** | **.036** |
| Age group x Main Effect | -.871 | .309 | -.582 | **.007** | **.028** |
| **OPA – R. middle occipital gyrus** | | | | | |
| Age Group | -.205 | .057 | -.458 | **<.001** | **.005** |
| Main effect | -.210 | .188 | -.142 | .269 | .296 |

Note: *b* = unstandardized coefficient; SE *b* = standard error of the unstandardized coefficient; β = standardized coefficient; FG = frontal gyrus; FS = frontal sulcus. Significant effects ($p < .05$) are in bold.

**Supplementary Table 5:** Results of the multiple regression analyses predicting source memory performance from age group and PPI main effects (seed-target connectivity change) during object encoding.

| Seed -Target | *b* | SE *b* | *β* | *p* | $p_{corrected}$ |
|---|---|---|---|---|---|
| **LOC - L. middle occipital gyrus** | | | | | |
| Age Group | -.228 | .058 | -.510 | **<.001** | **.002** |
| Main effect | -.093 | .112 | -.107 | .410 | .820 |
| **LOC - R. lateral occipital gyrus** | | | | | |
| Age Group | -.216 | .056 | -.483 | **<.001** | **.002** |
| Main effect | -.067 | .107 | -.079 | .531 | .820 |

Note: *b* = unstandardized coefficient; SE *b* = standard error of the unstandardized coefficient; β = standardized coefficient.

**Supplementary Table 6.** Results of the multiple regression analyses predicting source memory performance from age group and PPI main effects (seed-target connectivity change) during scene retrieval.

| Seed -Target | *b* | SE *b* | *β* | *p* | $p_{corrected}$ |
|---|---|---|---|---|---|
| **MPA - L. middle FG** | | | | | |
| Age Group | -.143 | .072 | -.320 | **.047*** | **.047** |
| Main effect | .584 | .166 | .753 | **<.001** | **.003** |
| Age Group x Main Effect | -.591 | .205 | -.741 | **.006** | **.018** |
| **MPA - L. intraparietal sulcus** | | | | | |
| Age Group | -.233 | .055 | -.521 | **< .001** | **.003** |
| Main Effect | .197 | .101 | .240 | .057 | .114 |
| **MPA - L. medial superior FG** | | | | | |
| Age Group | -.210 | .062 | -.468 | **.001** | **.003** |
| Main Effect | -.032 | .111 | -.040 | .776 | .776 |
| **PPA - L. middle FG** | | | | | |
| Age Group | -.230 | .057 | -.514 | **<.001** | **.002** |
| Main Effect | .117 | .092 | .160 | .211 | .422 |
| **PPA - L. inferior FG** | | | | | |
| Age Group | -.217 | .056 | -.485 | **<.001** | **.002** |
| Main Effect | .030 | .071 | .053 | .677 | .677 |

Note: *b*: unstandardized coefficient; SE *b*: standard error of the unstandardized coefficient; β: standardized coefficient; FG = frontal gyrus.

**Supplementary Table 7.** Results of the multiple regression analyses predicting source memory performance from age group and PPI main effects (seed-target connectivity change) during object retrieval.

| Seed-Target | *b* | SE *b* | *β* | *p* | $p_{corrected}$ |
|---|---|---|---|---|---|
| **L. LOC – L. inferior FG** | | | | | |
| Age Group | -.215 | .057 | -.481 | **<.001** | .006 |
| Main effect | -.041 | .158 | -.033 | .795 | 1.000 |
| **L. LOC – L. intraparietal sulcus** | | | | | |
| Age Group | -.218 | .057 | -.487 | **<.001** | .006 |
| Main effect | .017 | .162 | .014 | .916 | 1.000 |
| **L. LOC – L. supplementary motor cortex** | | | | | |
| Age Group | -.218 | .055 | -.487 | **<.001** | .006 |
| Main effect | -.246 | .142 | -.212 | .089 | .445 |
| **L. LOC – L. precuneus** | | | | | |
| Age Group | -.215 | .056 | -.481 | **<.001** | .006 |
| Main effect | -.104 | .190 | -.069 | .588 | 1.000 |
| **L. LOC – L. middle FG** | | | | | |
| Age Group | -.284 | .064 | -.634 | **<.001** | .006 |
| Main effect | -.812 | .364 | -.446 | **.031** | .186 |
| Age Group x Main Effect | .938 | .458 | .442 | **.046** | .276 |
| **L. LOC – R. cerebellum** | | | | | |
| Age Group | -.215 | .057 | -.479 | **<.001** | .006 |
| Main effect | -.061 | .210 | -.037 | .774 | 1.000 |

Note: *b*: unstandardized coefficient; SE *b*: standard error of the unstandardized coefficient; β: standardized coefficient; FG = frontal gyrus.
